# Cellular and subcellular specialization enables biology-constrained deep learning

**DOI:** 10.1101/2025.05.22.655599

**Authors:** Alessandro R. Galloni, Ajay Peddada, Yash Chennawar, Aaron D. Milstein

## Abstract

Learning and memory in the brain depend on changes in the strengths of synaptic connections between neurons. While the molecular and cellular mechanisms of synaptic plasticity have been extensively studied experimentally, much of our understanding of how plasticity is organized across populations of neurons during task learning comes from training artificial neural networks (ANNs) using computational methods. However, the architectures of modern ANNs and the algorithms used to train them are not compatible with fundamental principles of neuroscience, leaving a gap in understanding how the brain coordinates learning across multiple layers of neural circuitry. Here we leverage recent experimental evidence to test an emergent theory that biological learning depends on specialization of distinct neuronal cell types and compartmentalized signaling within neuronal dendrites. We demonstrate that multilayer ANNs comprised of separate recurrently connected excitatory and inhibitory cell types, and neuronal units with separate soma and dendrite compartments, can be trained to accurately classify images using a fully biology-compatible deep learning algorithm called *dendritic target propagation*. By adhering to strict biological constraints, this model is able to provide unique insight into the biological mechanisms of learning and to make experimentally testable predictions regarding the roles of specific neuronal cell types in coordinating learning across different brain regions.

## Main

When an animal learns to perform a task, patterns of neuronal activity in the brain change over time as behavioral performance improves^1^. Mathematically, this learning can be described as an optimization process that minimizes task performance error by modifying the parameters in the brain that control behavior. Thus, by definition, when learning is successful, neuronal parameters must have changed in a direction that minimizes performance error^2^. However, it is not known how the brain utilizes information about behavioral outcomes to adjust the right parameters in the right neurons during learning – the credit assignment problem^3^ – or how it transmits that information to coordinate learning across brain areas or multiple layers of neural circuitry^4^.

In standard models of learning in neural circuits, the primary parameters that can be adjusted during training are the strengths of synaptic connections between neuronal units, often referred to as “synaptic weights.” In the machine learning field, synaptic weights in artificial neural networks (ANNs) are typically optimized using gradient descent. This requires calculating the impact of each individual synapse on the overall performance error of the network (the error gradient with respect to the network weights), and precisely adjusting each synaptic weight proportionally. In multilayer ANNs, an algorithm called “backpropagation” is typically used to transmit gradient information across multiple neural circuit layers^5,6^. However, it has not been demonstrated how backpropagation could be implemented using entirely biology-compatible mechanisms^7^. In contrast, in the neuroscience field, synaptic weights in neural circuit models are typically modified using Hebbian synaptic plasticity rules that have been observed experimentally, like spike-timing-dependent plasticity (STDP)^8-10^, but those rules do not necessarily adjust synaptic weights in the direction of the error gradient or guarantee improvement in task performance^4,11^. In this study, we aim to reconcile the discrepancies between these two approaches and to demonstrate a mechanism for performance error to guide learning in multilayer neural circuits in a way that is compatible with known principles of neurobiology.

In particular, we impose the following experimentally-supported biological constraints to address multiple biologically-implausible features of standard ANNs and the backpropagation algorithm: 1) the *cellular diversity* constraint: while neuronal units in standard ANNs are uniform, biological neurons are specialized into diverse cell types with different functions^12^ (Fig. 1); 2) the *signed synapse* constraint: while gradient descent can cause a synaptic weight to change sign (from positive to negative, or vice versa), biological synapses cannot change sign, and neurons are typically pre-specified during development as either excitatory (all outgoing synapses are positive) or inhibitory (all outgoing synapses are negative) (this constraint is often referred to as Dale’s law^13^); 3) the *continuous signaling* constraint: while backpropagation requires that sensory inference and error propagation occur in two separate phases (sensory responses flow bottom-up exclusively during the sensory phase, and error signals flow top-down exclusively during the learning phase), in biological circuits, neurons continuously send and receive both bottom-up and top-down signals; and 4) *the non-symmetric synapse* constraint: while backpropagation requires that the synaptic weights used to propagate sensory information bottom-up are exactly symmetrical to those used to propagate gradient signals top-down, in biological circuits separate synapses with distinct weights are responsible for processing either bottom-up or top-down signals (this is often referred to as the “weight transport problem”^14-16^).

**Figure 1.**
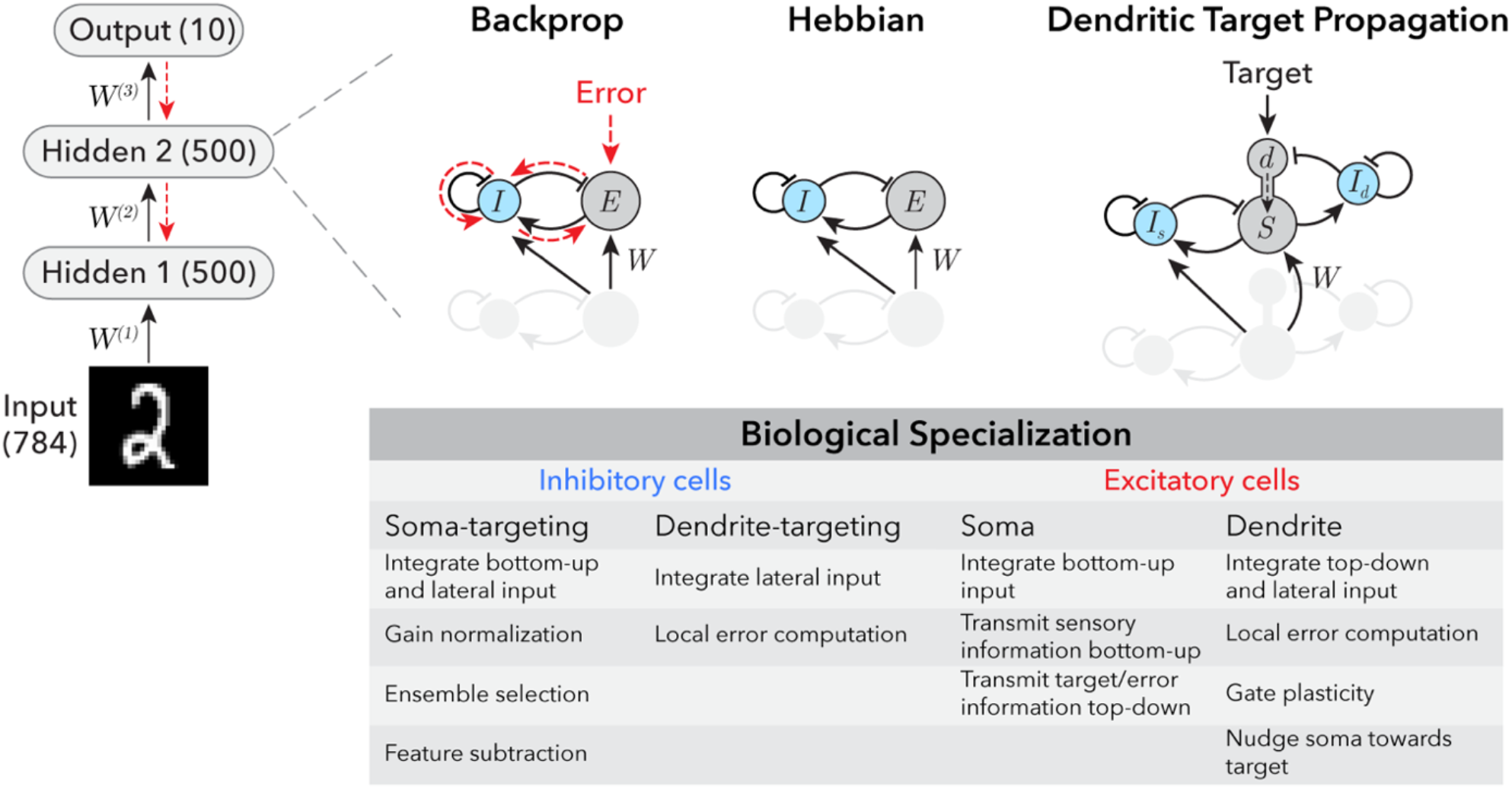
Incorporating biological cellular and subcellular specialization into artificial neural networks. *Top left:* Architecture of multi-layer neural networks trained to classify images of handwritten digits. The number of excitatory units in each layer is indicated in parentheses. *Top right:* Each excitatory-inhibitory artificial neural network (EIANN) has two hidden layers containing 500 excitatory (E) units and 50 inhibitory (I) units that are recurrently connected, consistent with experimentally observed proportions. Networks trained with the proposed biology-constrained algorithm *dendritic target propagation* have two separate populations of 50 inhibitory neurons each that target either the soma (*I*_*s*_) or the dendrite (*I*_*d*_). Networks trained with backpropagation receive top-down error signals, Hebbian networks receive no top-down inputs, and networks trained with *dendritic target propagation* receive top-down target signals and compute error locally. *Bottom*: Table summarizes proposed computational roles of different biological cell types.

Finally, we propose a biologically-constrained mechanism for information about performance error to be computed and propagated within a multilayer neural circuit. Our approach depends on the morphology of biological neurons, which contain a tree-like structure of branched subcellular compartments called dendrites. We leverage insight from recent experimental and theoretical work supporting specialized roles for electrically compartmentalized dendrites in processing top-down signals and controlling synaptic plasticity^17-34^. The algorithm, which we refer to as *dendritic target propagation*, locally computes an error signal in the distal dendrites of excitatory neurons as a difference between top-down excitation and lateral feedback inhibition^18,19,35^ (Fig. 1). In each neuron, this dendritic error signal has two effects: it determines the sign and magnitude of synaptic weight updates, and it “nudges” the activity level of the neuron either up or down. These changes in activity are then relayed to lower layers of the network via top-down projections.

This, in turn, signals to neurons in lower layers that they must also adjust their synaptic weights to reduce network performance error. We demonstrate that *dendritic target propagation* can be used to effectively train multilayer neural networks with biologically-constrained recurrent architectures to perform stimulus classification tasks. We also show that this algorithm is compatible with multiple experimentally and theoretically supported synaptic plasticity rules, and remains effective even when the weights of bottom-up and top-down synapses are independently initialized and learned. In summary, we demonstrate that neurobiological principles like cell-type diversity, subcellular computation in dendrites, and synaptic plasticity can support target-directed deep learning in the brain.

### Training recurrent networks with separate excitatory and inhibitory neurons

First, we imposed the biological *cellular diversity* and *signed synapse* constraints on the architecture of a 3-layer ANN (Fig. 1) by dividing neurons within each layer into separate excitatory (E) and inhibitory (I) populations. Within each layer, both excitatory and inhibitory cells receive bottom-up inputs from excitatory neurons in the previous layer, and lateral inputs from inhibitory neurons in the same layer. This establishes a feedforward inhibitory circuit motif common in mammalian hippocampal and cortical circuits, which has been associated with specific computations such as input normalization and stimulus feature subtraction^36,37^ (Fig. 1). Inhibitory neurons also receive a lateral input from excitatory neurons in the same layer. This establishes a recurrent feedback inhibitory circuit motif, which can impose competition between ensembles of excitatory neurons, limit the number of neurons that can be simultaneously active (i.e. enforce sparsity), shape their stimulus selectivity, and decorrelate their activity^38-40^ (Fig. 1).

Despite potential advantages of this functional division-of-labor across neuronal cell types (Fig. 1), this biological microcircuit architecture imposes some additional challenges. Depending on the strength of connections and timescales of input integration, recurrent excitatory-inhibitory (EI) networks can be prone to unstable or oscillatory dynamics^41-43^, which can interfere with learning^44^. Furthermore, biological neuronal circuits are typically comprised of a majority of excitatory neurons and a minority of inhibitory neurons^45,46^, potentially imposing a bottleneck for memory storage^47^ and computations that require subtraction^48,49^. Thus, before testing biological learning mechanisms, we first verified that the backpropagation algorithm can be used to train a multilayer excitatory-inhibitory ANN (EIANN) to perform an image classification task (handwritten digit characters) (Fig. 1; Methods). We tuned model hyperparameters to ensure that dynamic network responses to static image stimuli converged to a non-oscillatory equilibrium (Supplementary Fig. S1; Methods). The resulting trained EIANN exhibited classification accuracy comparable to a standard feedforward network (Fig. 2a,b,g). We used an activity maximization method to visualize the stimulus preferences of individual neurons after training^50^, and found that excitatory units in the EIANN had more spatially structured receptive fields than the non-sign-constrained units in the standard ANN (Fig. 2d,e,i; Methods)^51^. Importantly, these receptive fields contained some regions of the stimulus space that were excitatory (Fig. 2d,e; black), and some that were inhibitory (Fig. 2d,e; blue), indicating that separating excitation and inhibition into distinct neuronal cell types did not interfere with the capability of the network to perform stimulus feature subtraction – an important elementary computation. Importantly, training a network without any inhibitory neurons (all positive weights) resulted in strongly reduced performance (Supplementary Table S1), indicating that the task requires a source of negative weights to perform stimulus feature subtraction^52^.

**Figure 2.**
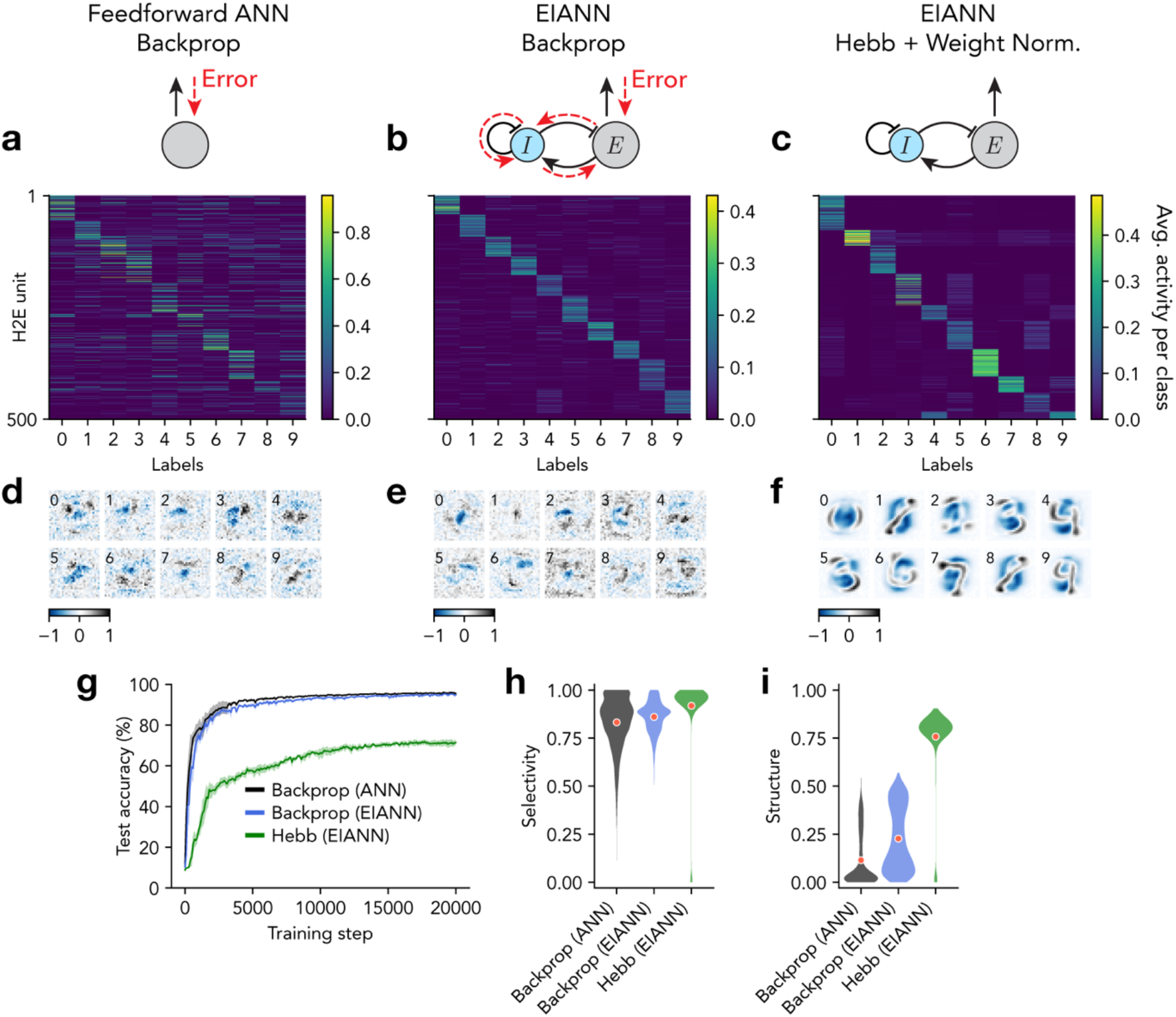
Coordinating learning across network layers requires top-down signaling. **a-c**, Average neuronal population activity in response to each class of handwritten digits. **a**, Shown are 500 excitatory (E) units in the second hidden layer (H2) of a feedforward ANN trained with backpropagation. **b**, Same as **a** for a recurrent EIANN trained with backpropagation. **c**, Same as **a** for a recurrent EIANN trained with a normalized Hebbian rule. **d-f**, Example receptive fields selected from H2E units from the networks shown in **a-c**. Numbers in the top left corners indicate the label of the class that produces the largest activity (averaged across samples) for each unit. **g**, Classification performance accuracy. Shading indicates standard deviation across five instances of each network. **h**, Selectivity of E units (in both hidden layers) over stimulus classes. **i**, Spatial structure of the receptive fields of E units (in both hidden layers), as measured by spatial autocorrelation (Moran’s I).

Following training, excitatory neurons exhibited a greater degree of stimulus selectivity than inhibitory neurons (Fig. 2a,b,h and Supplementary Fig. S2), consistent with experimental recordings from the mammalian hippocampus and cortex ^53-57^. Interestingly, we found a similar degree of stimulus selectivity in inhibitory cells when all incoming and outgoing synapses to and from inhibitory cells were fixed at their random initial weights and not learned (Supplementary Fig. S2). Fixed inhibition also resulted in comparable classification performance, suggesting that plasticity of inhibition is not required for the task (Supplementary Fig. S2).

Next, we sought to impose the biological *continuous signaling* constraint, which disallows neurons from switching between separate signaling modes in which their activities represent either bottom-up responses to the sensory environment or top-down error gradients. The simplest implementation of this constraint is to completely remove top-down connections between circuit layers, and to use unsupervised learning rules to promote self-organization of each layer such that they produce discriminable patterns of activity in response to different classes of stimuli^58-61^. Then, an error-dependent learning rule can be used in the top layer to perform image classification without propagating error signals to earlier network layers. A recent study^62^ used a combination of experimentally-supported Hebbian and homeostatic learning rules^63-65^ at all synapses within a single layer of an EIANN to achieve highly selective unit responses and highly discriminating population responses that uniformly tiled a stimulus space, without any supervision. In that network, feedback inhibition imposed competition between units (Fig. 1), encouraging different units to represent different stimuli. At each synapse between a presynaptic neuron *j* and a postsynaptic neuron *i*, the connection weight *w*_*ij*_ was updated proportional to the correlation *c*_*ij*_ of the activities of the two units:

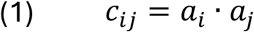

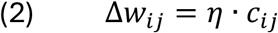

where *η* is a learning rate. After each weight update, the total pool of synaptic resources available in each postsynaptic neuron for a specific projection type (e.g. bottom-up excitatory inputs) was maintained constant by applying the following weight normalization inspired by experimentally-observed heterosynaptic plasticity^62,63,65^:

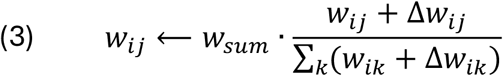

where *w*_*sum*_ is a scale factor. This normalization prevents unbounded growth of synaptic strengths, and enhances stimulus selectivity by imposing competition between synapses within a neuron^62^. We implemented this unsupervised Hebbian learning rule with weight normalization in our multilayer EIANN (Fig. 1 and Fig. 2c; Methods) and found that, while hidden units became highly stimulus-selective after learning (Fig. 2c,h), their receptive fields showed little diversity, resembling the average image within a class (Fig. 2f and Supplementary Fig. S3). Some neurons responded similarly to more than one class of stimuli (Fig. 2c and Supplementary Fig. S3), resulting in poor image classification performance (Fig. 2g). These results suggest that, while unsupervised Hebbian learning rules are effective at self-organizing neuronal EI networks to produce sparse and selective sensory responses, without top-down signaling between layers in a multilayer network, a purely unsupervised algorithm is unable to coordinate learning across layers or to guarantee that the learned stimulus representations will be useful for accurate image classification (Supplementary Fig. S3).

### Biology-constrained multilayer credit assignment via dendritic target propagation

The above results suggested that top-down propagation of error information between circuit layers is essential for target-directed learning. However, the biological *continuous signaling* constraint dictates that neurons simultaneously propagate their activity both bottom-up and top-down, which means that activity generated in higher layers could interfere with sensory responses in lower layers, and likewise sensory information could be aliased with information about learning targets or performance error. Previous models have proposed various solutions to this problem, including i) limiting the strengths of top-down connections^66^, ii) applying different temporal filters to bottom-up and top-down pathways^24,26,27^, iii) processing bottom-up and top-down signals in separate neurons^67-69^, iv) processing them in separate subcellular compartments (soma and dendrite)^21-24,70^, and/or v) limiting the influence of top-down inputs by predicting and cancelling them with a separate lateral or top-down input pathway^21,23,27^. Only recently has experimental data from awake behaving animals become available that tests some of the predictions of these models and provides insight into the roles of top-down excitation and lateral inhibition to the distal apical dendrites of pyramidal neurons in modulating neuronal activity and plasticity during target-directed learning^17-19,31,35,71-76^.

We sought to incorporate into our EIANN model the following experimental findings from the mammalian hippocampus and cortex: 1) sufficient input to the apical dendrites of excitatory pyramidal cells can evoke a long (∼50 ms to > 1 s) regenerative event called a dendritic calcium spike^29,77-80^; 2) dendritic calcium spikes can propagate to the soma, resulting in an increase in output firing rate; 3) dendritic calcium spikes can rapidly induce either increases or decreases in synaptic strength through a potent form of synaptic plasticity called behavioral timescale synaptic plasticity (BTSP)^17-19,29,31^, 4) top-down excitatory inputs onto distal apical dendrites carry information about behaviorally-relevant context, cues, and goals, and strongly promote dendritic calcium spikes and associated plasticity^19,31,71,76^, and 5) dendritic inhibition from local interneurons strongly inhibits dendritic calcium spikes and associated plasticity^35,57,74,75,81^. Accordingly, we modified the excitatory neurons in our EIANN to comprise separate soma and distal apical dendrite compartments (Fig. 1 and Fig. 3a, Methods). Bottom-up excitatory inputs and lateral inhibitory inputs from soma-targeting interneurons arrive at the soma compartment (representing the cell body as well as perisomatic dendrites) (Fig. 1). We also added an additional specialized population of dendrite-targeting interneurons, which receive lateral excitatory inputs and recurrent inhibitory inputs from each other (Fig. 1 and Fig. 3a). The apical dendrite compartment of excitatory neurons receives top-down excitatory inputs, and lateral inhibitory inputs from the dendrite-targeting interneurons (Fig. 1 and Fig. 3a). To train this dendritic EIANN, we designed a learning algorithm termed *dendritic target propagation*, in which the activation state of the dendritic compartments of excitatory neurons, *D*, acts both as a gate for plasticity and as a source of input to the soma compartments that affects their output firing rates.

**Figure 3.**
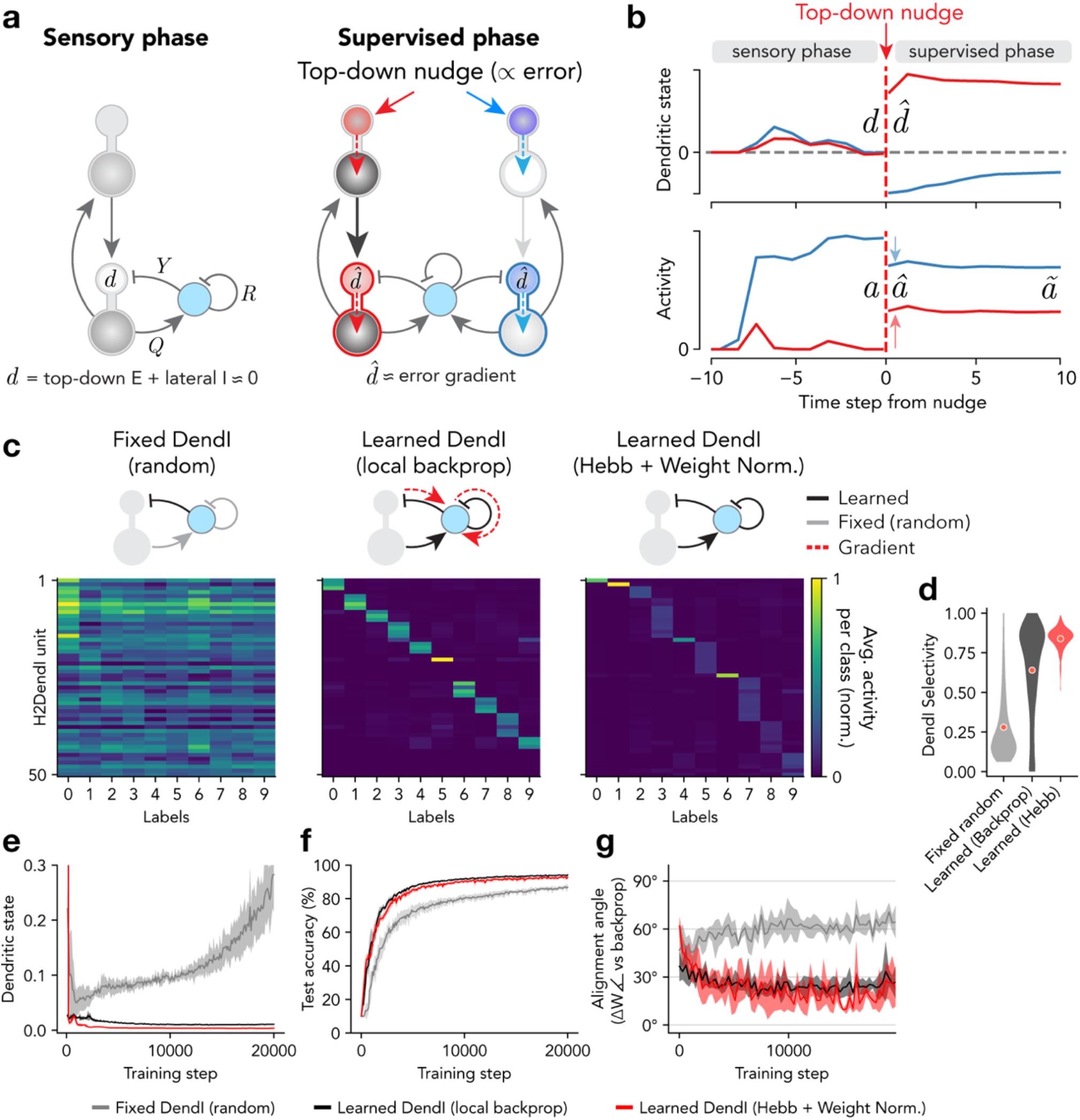
Adaptation of dendrite-targeting inhibition improves deep credit assignment. **a**, Diagram of the *dendritic target propagation* algorithm. *Left:* Network responses to sensory stimuli are propagated both forward and backward through the network layers. During this sensory phase, lateral dendritic inhibition aims to accurately cancel top-down dendritic excitation, resulting in dendritic states close to zero. *Right:* During a supervisory phase, output layer excitatory units are nudged up (red) or down (blue) to their target activity. These changes in activity are propagated to lower layer dendrites, where they are unpredicted and remain uncancelled by dendritic inhibition, resulting in dendritic states that approximate local error gradients in each neuron. **b**, Example temporal dynamics of the dendritic states (top) and resulting activities (bottom) of the two example hidden layer neurons illustrated in **a**. The caret symbol (^) represents values immediately after the nudge is applied, while the tilde symbol (∼) represents values after additional time steps of equilibration. **c**, Average neuronal population activity in response to each class of handwritten digits are shown for dendrite-targeting inhibitory neurons (DendI) in the second hidden layer (H2) for networks using three different learning schemes for their incoming weights. *Left:* Weights onto DendI neurons are frozen at their random initialization. *Middle:* Weights onto DendI neurons are learned by using backpropagation locally within each hidden layer to minimize the magnitude of excitatory neuron dendritic states. *Right:* Weights onto DendI neurons are learned using a fully unsupervised normalized Hebbian learning rule. **d**, Selectivity of DendI units (in both hidden layers) over stimulus classes. **e**, Average absolute value of the sensory-phase dendritic states of hidden layer E cells during training. **f**, Classification performance accuracy. **g**, For each model, the bottom-up excitatory (*W*) weight updates prescribed by the model were compared to those prescribed by backpropagation using cosine similarity. An alignment angle of 90° indicates an update that is orthogonal to the gradient, while 0° reflects identical weight updates. **e-g**, Shading indicates standard deviation across five instances of each network.

In a network with *L* layers, the dendritic states in the output layer *D*^(*L*)^ are initially set to zero during stimulus presentation, in the absence of any supervisory input. However, any activity *A*^(*E,L*)^ in output layer excitatory neurons excites the dendrites of neurons in the previous hidden layer *l* = *L* − 1 via top-down projections with weights *B*^(*L*)^:

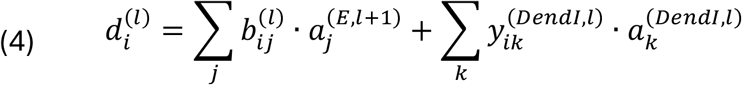

where 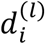 is the dendritic state of a postsynaptic excitatory neuron *i* in layer *l*, 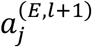 is the activity of a presynaptic excitatory neuron j in layer *l* + 1 connected to 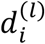 via weight 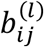, and 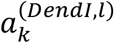 is the activity of a presynaptic dendrite-targeting inhibitory neuron *k* in layer *l* connected to 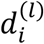 via weight 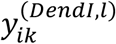 (note that here we use capitalized symbols to indicate population activity vectors and weight matrices, and lowercase symbols to indicate single scalar values for one cell, compartment, or synapse). We initially set the top-down weights *B*^(*l*)^ (from layer *l* + 1 to layer *l*) as symmetric to the bottom-up weights *W*^(*l* +1)^ (from layer *l* to layer *l* + 1), as prescribed by the backpropagation algorithm. Later we will consider methods to separately learn the independent weights *B*^(*l*)^.

In order for the firing rates of neurons to cleanly reflect their sensory responses without interference from top-down inputs, ideally the dendritic states *D*^(*l*)^ in each hidden layer *l* would also be close to zero during stimulus presentation (Fig. 3a,b). Within the dendritic EIANN architecture, this can be achieved if the sum of lateral inhibitory inputs to each dendrite exactly predicts and cancels the sum of its top-down excitatory inputs (Equation (4)). Therefore, we applied a learning rule to the lateral inhibitory synapses onto excitatory neuron dendrites (with weights *Y*^(*l*)^) that aims to minimize the dendritic states *D*^(*l*)^:

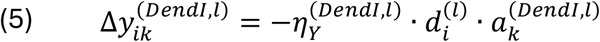

for postsynaptic excitatory neuron *i* and presynaptic dendrite-targeting inhibitory neuron *k* in layer *l*, where 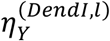 is a learning rate.

Following equilibration of neuronal activities during the sensory phase, we then applied supervisory inputs *N* to the dendrite compartments *D*^(*L*)^ of excitatory neurons in the output layer to “nudge” their activity *A*^(*E,L*)^ towards target values *T* that would lead to correct stimulus categorization (Fig. 3a,b):

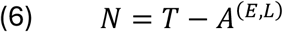

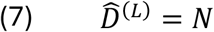

where the caret symbol (^) indicates variables after the supervisory inputs have been applied. In each layer *l*, the “nudged” dendritic states 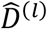 in turn “nudge” the excitatory neuron firing rates *Â*^(*E,l*)^ (Fig. 3a):

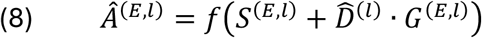

where *S*^(*E,l*)^ is the state of the soma compartments of the excitatory neurons, *f* is the rectified linear unit activation function (ReLU) (see Methods), and *G*^(*E,l*)^ = *f* ^′^ *ES*^(*E,l*)^H is the derivative of the ReLU activation function, which is one when *S*^(*E,l*)^ > 0 and zero otherwise. This change in output layer activity then propagates to lower layers via top-down projections (see Methods). Since lateral dendritic inhibition is tuned to cancel the hidden layer dendritic states *D*^(*l*)^ during the sensory phase, these changes in top-down excitation during the supervisory phase are not predicted by the inhibitory neurons and remain uncancelled (Fig. 3a,b). This results in the “nudged” dendritic states 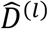 in each layer reflecting a weighted sum of the supervisory nudges 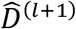applied to the layer above. Finally, the weights *W*^(*l*)^ of the bottom-up excitatory inputs onto layer *l* excitatory neurons are updated with the following dendritic state-dependent learning rule:

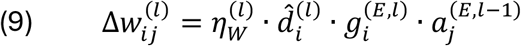

for postsynaptic excitatory unit *i* in layer *l* and presynaptic excitatory unit *j* in layer *l* − 1. To the extent that 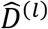 approaches the value of the error gradient 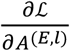 used in backpropagation (see Methods), this weight update will also move the network in a direction that minimizes error.

The above scheme locally computes error-related learning signals in the dendrites of excitatory neurons (Fig. 3a). However, it does not yet address whether or how synaptic weights should be updated at the inputs to dendrite-targeting interneurons to ensure that their activity provides an appropriate basis for cancelling the top-down excitatory inputs to the excitatory neuron dendrites. These interneurons receive lateral excitatory inputs with weights *Q*^(*DendI,l*)^ and recurrent inhibitory inputs with weights *R*^(*DendI,l*)^ (Fig. 3a). First, we asked whether the *dendritic target propagation* algorithm could effectively train the dendritic EIANN if the weights of these projections were simply held fixed at their random initial values, and not learned, as proved effective for soma-targeting inhibitory neurons (Supplementary Fig. S2). Fixed dendritic inhibition resulted in low stimulus selectivity in dendrite-targeting interneurons (Fig. 3c,d), ineffective cancelling of top-down sensory input to excitatory neuron dendrites (Fig. 3e), and poor image classification performance by the network (Fig. 3f). Updates to the bottom-up excitatory weights *W*^(*l*)^ were poorly aligned with the direction of the error gradient (Fig. 3g). This implies that, unlike in soma-targeting interneurons, plasticity in dendrite-targeting interneurons may be required to improve local error computation and task performance.

Before testing biologically-plausible learning mechanisms in these dendrite-targeting interneurons, we first took a normative approach and tested whether task performance could be improved if the weights of inputs onto dendrite-targeting inhibitory neurons were directly optimized by gradient descent to minimize the dendritic states of the excitatory neurons. To do this, we defined a separate loss function ℒ^(*l*)^ in each hidden layer as the mean squared dendritic state of the excitatory neurons in the layer:

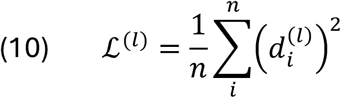

Then, in each hidden layer, we used backpropagation to adjust weights in proportion to the gradient of this local loss function with respect to the synaptic weights *Q*^(*DendI,l*)^ and *R*^(*DendI,l*)^ (see Methods). Learning dendritic inhibition with backpropagation resulted in dendrite-targeting interneurons that were highly stimulus selective (Fig. 3c,d), excitatory dendritic states that were effectively minimized (Fig. 3e), and increased task performance (Fig. 3f).

With this configuration of dendritic inhibition learned by gradient descent, the *dendritic target propagation* algorithm prescribed updates to the bottom-up excitatory weights *W*^(*l*)^ that were well-aligned with those prescribed by the backpropagation algorithm (Fig. 3g). Given this gradient-based baseline to compare against, we next explored how the weights of inputs to the dendrite-targeting interneurons could be learned with biologically-plausible learning rules. To address this, we revisited the normalized Hebbian plasticity rule previously used to self-organize individual circuit layers without supervisory inputs^62^ (Fig. 1 and Fig. 2c; see Methods). This approach has been shown to be directly related to principal component analysis (PCA) and independent component analysis (ICA), and can enable a population of neurons to learn a compressed (reduced dimensionality) representation of their inputs ^62,82^. We found that applying the normalized Hebbian rule to the incoming projections onto dendrite-targeting interneurons resulted in high stimulus selectivity (Fig. 3c,d), effective minimization of excitatory dendritic states (Fig. 3e), increased classification performance (Fig. 3f and Fig. 4b), and bottom-up weight updates that better aligned with backpropagation (Fig. 3g). This strategy also resulted in highly selective hidden layer excitatory neurons (Fig. 4a,d) with highly structured receptive fields (Fig. 4c,d and Supplementary Fig. S3). These results demonstrate that unsupervised Hebbian learning in inhibitory neurons can be combined with dendritic-state-dependent learning in excitatory neurons to achieve biology-compatible training of multilayer networks. They also highlight that while unsupervised Hebbian rules are limited in the complete absence of top-down supervision (Fig. 1 and Fig. 2c), they can be used to effectively self-organize some components of a neural circuit, as long as they are supplemented by top-down modulation of learning in other components of the circuit, which can provide a scaffold to organize around.

**Figure 4.**
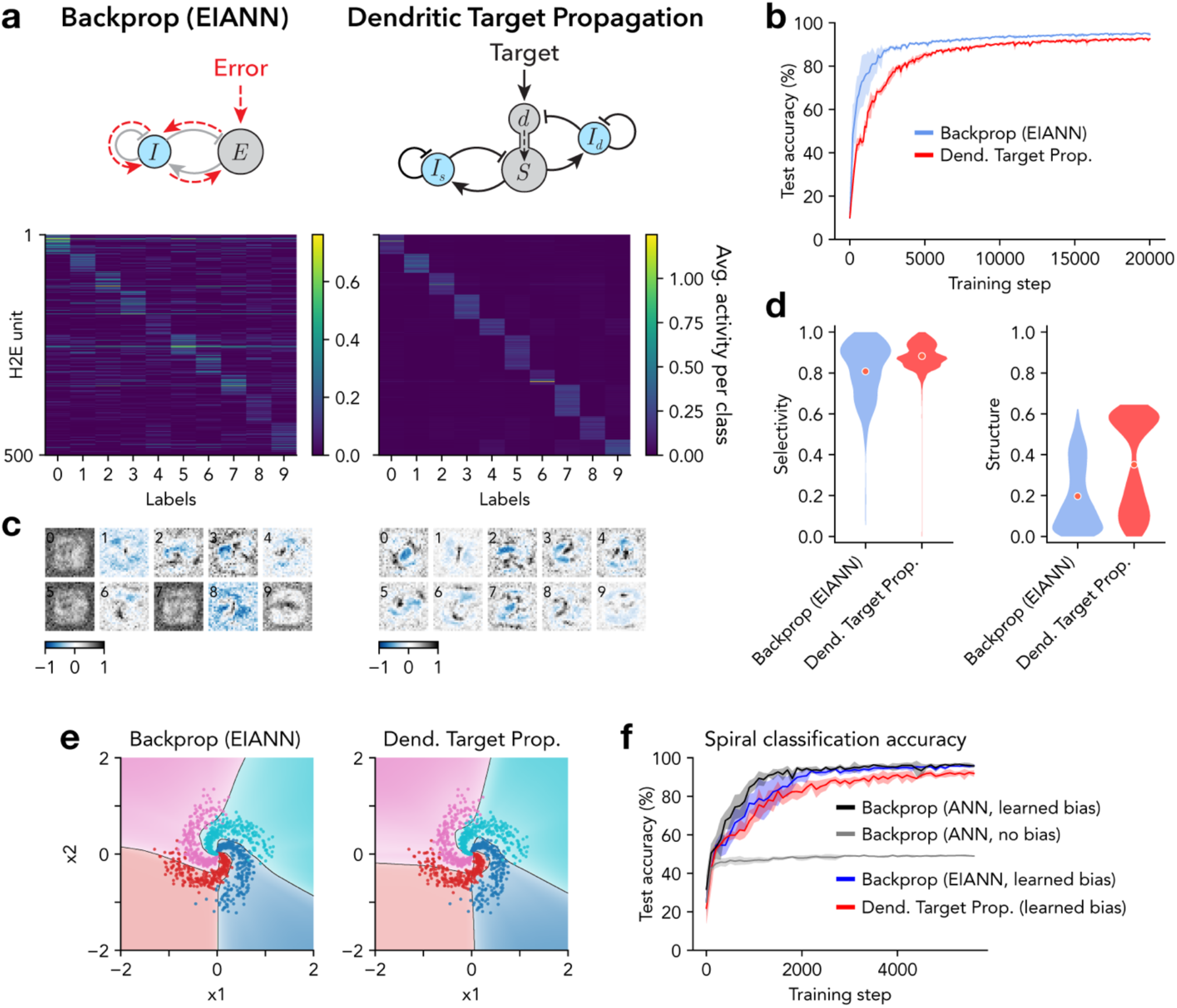
Dendritic target propagation enables biology-constrained deep learning. **a**, Average neuronal population activity in response to each class of handwritten digits. *Left*: Shown are 500 excitatory (E) units in the second hidden layer (H2) of a recurrent EIANN trained with backpropagation. *Right*: Same as *left* for a dendritic EIANN trained with *dendritic target propagation*. **b**, Classification performance accuracy. **c**, Example receptive fields selected from H2E units from the networks shown in **a**. Numbers in the top left corners indicate the label of the class that produces the largest activity (averaged across samples) for each unit. **d**, *Left*: Selectivity of E units (in both hidden layers) over stimulus classes. *Right*: Spatial structure of the receptive fields of E units (in both hidden layers), as measured by spatial autocorrelation (Moran’s I). **e**, *Left*: Following training on the 2D spirals classification task, nonlinear decision boundaries are shown for an EIANN trained with backpropagation. Data points with colors that do not match the background region were misclassified. *Right*: Same as *left* for a dendritic EIANN trained with *dendritic target propagation*. **f**, Classification performance accuracy on the 2D spirals task. The networks shown in **e** are compared to standard feedforward ANNs with and without learned neuronal bias parameters. In **b** and **f**, shading indicates standard deviation across five instances of each network.

For simplicity, the models described above were all trained by adjusting synaptic connection weights, but did not include learnable neuronal bias parameters. Biases are typically implemented as modifiable levels of positive or negative background input to each neuronal unit that affect how much additional input a cell needs to cross its activation threshold:

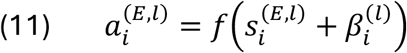

Some nonlinear classification tasks cannot be solved without adjustable biases. Thus, we tested the effectiveness of the *dendritic target propagation* algorithm on a 2D spiral classification task that requires learned biases (Supplementary Table S2). We added neuronal bias parameters 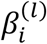 to excitatory neurons only, and updated them with a learning rule that depends linearly on the dendritic state:

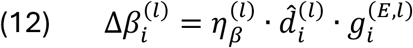

The resulting nonlinear classification boundaries are visualized in Fig. 4e. This method resulted in classification accuracy comparable to backpropagation (Fig. 4f and Supplementary Table S2), demonstrating that the *dendritic target propagation* algorithm is also applicable to highly nonlinear classification tasks that have a strong dependence on learned biases.

### Dendritic target propagation is compatible with multiple biological synaptic plasticity rules

In the above section, we described a biology-constrained method that performs deep synaptic credit assignment by computing local error signals in the activation states of the distal dendrites of excitatory neurons. These dendritic error signals were used both 1) to “nudge” the activity of excitatory neurons towards target values that would reduce task performance error, and 2) to update bottom-up synaptic weights with a learning rule that depended linearly on the dendritic state (linear dendritic state rule; LDS; Equation (9)). This scheme was inspired by the recently experimentally-observed BTSP rule, which depends on dendritic calcium spikes^17,18^. However, while the LDS rule was designed to directly approximate gradient descent, the BTSP rule has additional unique features not captured by the simple LDS rule, including saturable weights and the ability to modify inputs that are active at long time delays (up to ∼5 s) from a dendritic calcium spike^17,18^. Thus, we investigated whether the more complex BTSP rule could also be used to update synaptic weights once the dendritic states of excitatory neurons have been set by *dendritic target propagation*. We also considered an alternative possibility that, once dendritic error signals have been used to “nudge” somatic activities, classical Hebbian plasticity rules that depend on somatic firing rates could also be used to update synaptic weights^60,66,83-86^. For this purpose, we tested two variants of Hebbian learning that have been well-supported by experimental and theoretical evidence: a “temporally contrastive Hebbian” (TCH) rule^66,84^, which is related to experimentally-observed STDP^8-10^, and the Bienenstock-Cooper-Munro (BCM) rule^85^, which features an adaptive plasticity threshold and is commonly used to model learning with firing rate homeostasis, such as during visual system development^60,86^.

The TCH rule weakens connections with correlated activity during the sensory phase, and strengthens connections that are correlated during the supervised phase:

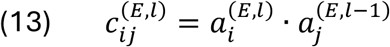

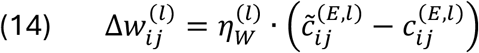

where the tilde symbol (∼) reflects variables after supervisory inputs have been applied and the network has re-equilibrated (Fig. 3a and Fig. 5b; see Methods). We expected this rule to perform similarly to the LDS rule, as TCH indirectly recovers the dendritic error signal by detecting how much somatic activities have changed after receiving the “nudge” from their dendrites:

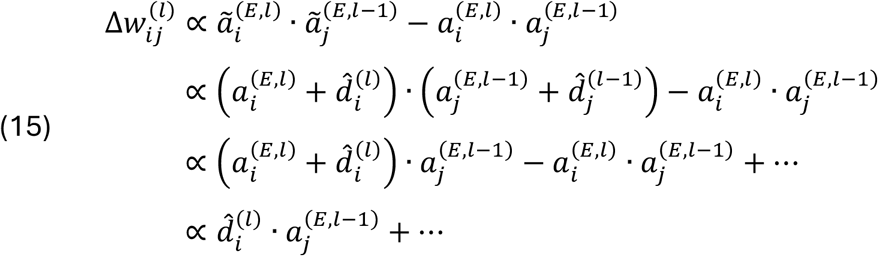

**Figure 5.**
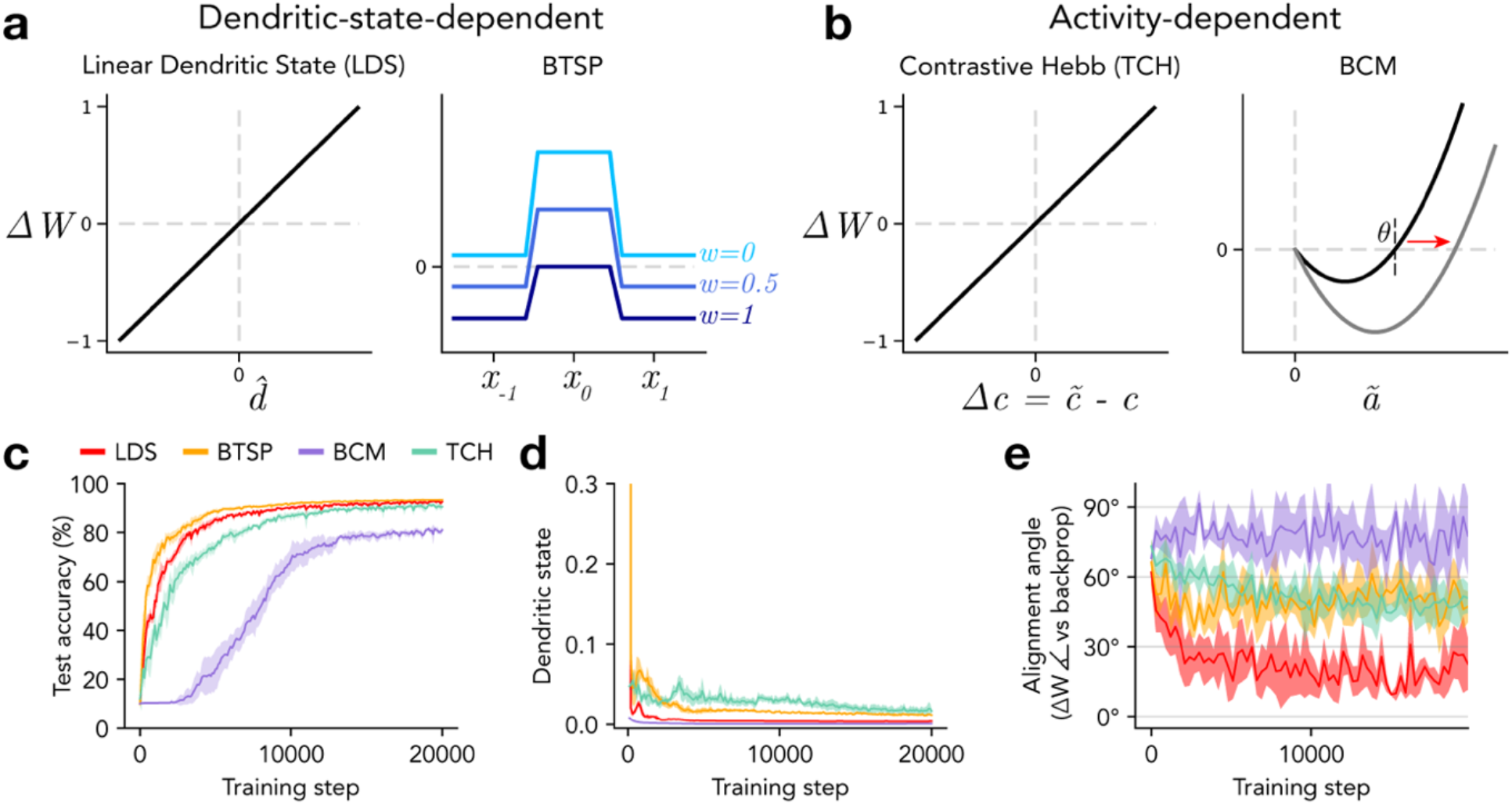
Dendritic target propagation is compatible with multiple biological synaptic plasticity rules. **a-b**, Diagrams summarize biological synaptic plasticity rules. **a**, Two plasticity rules that depend on dendritic state are represented. *Left:* The linear dendritic state (LDS) rule updates weights proportionally to dendritic state, and approximates backpropagation to the extent that dendritic state approximates the error gradient. *Right:* The behavioral timescale synaptic plasticity (BTSP) rule considers interactions between dendritic calcium spikes and presynaptic activity at long time delays, enabling interactions between temporally adjacent stimuli. BTSP also features a dependence on synaptic weight such that weak weights increase and strong weights decrease. **b**, Two plasticity rules that depend on somatic activity are represented. *Left:* The temporally contrastive Hebbian (TCH) rule updates weights proportionally to differences in activity correlations between a sensory phase and a supervised phase. *Right:* The BCM rule is a model of learning with firing rate homeostasis and an adaptive plasticity threshold that depends on the recent history of somatic activity. **c**, Classification performance accuracy. **d**, Average absolute value of the sensory-phase dendritic states of hidden layer E cells during training. **e**, Alignment angle between bottom-up excitatory (*W*) weight updates prescribed by each model and those prescribed by backpropagation. Shading indicates standard deviation across five instances of each network.

Indeed, we found that the TCH rule results in low amplitude dendritic states during the sensory phase (Fig. 5d) and high task performance comparable to the LDS rule (Fig. 5c), though it prescribed weight updates that were less aligned with backpropagation (Fig. 5e).

The BCM rule strengthens active connections when the postsynaptic activity is above an adaptive threshold *θ*^(*l*)^, and depresses them otherwise:

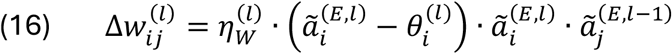

where the threshold 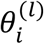 is a running average of activity 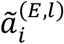 over recent stimulus samples

(Fig. 5b; see Methods). While this rule is calibrated to produce stable and selective receptive fields, it refers only to target-aligned activities but does not incorporate any form of error signals, even indirectly. Using this rule to train a dendritic EIANN resulted in very low dendritic states during the sensory phase (Fig. 5d), reduced image classification accuracy (Fig. 5c), and weight updates that were nearly orthogonal to gradient descent (Fig. 5e).

The BTSP rule is distinct in that it updates the weights of inputs that are active within a broad window spanning seconds before, during, and after a dendritic calcium spike^17,18^. It achieves this by referring to slowly decaying traces of recent presynaptic activity (eligibility traces *ET*^(*l*−1)^) and recent postsynaptic dendritic calcium spikes (instructive signals *IS*^(*l*)^)^18^:

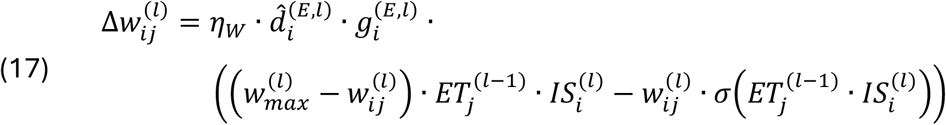

where 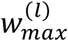 is a maximum weight and *σ* is a sigmoidal function (see Methods). This plasticity rule promotes selectivity by favoring potentiation of inputs activated by the current stimulus, and depressing inputs activated by temporally adjacent stimuli (Fig. 5a). When the BTSP rule was used to update the bottom-up weights of a dendritic EIANN trained with *dendritic target propagation*, it resulted in effective dendritic error computation (Fig. 5d) and task performance comparable to the LDS rule (Fig. 5c). This is particularly interesting given that the BTSP rule sometimes prescribes weight updates with opposite sign compared to the LDS rule and backpropagation, resulting in intermediate alignment between the two weight updates (Fig. 5e).

In summary, when the biology-constrained *dendritic target propagation* algorithm is used to locally compute error signals in the distal dendrites of excitatory neurons in a multilayer EIANN, a variety of experimentally-supported biological plasticity rules can flexibly be used to update the synaptic weights of bottom-up excitatory inputs. We found that synaptic plasticity rules like BTSP or TCH that refer either directly or indirectly to an error signal outperform rules like BCM that refer only to somatic activity.

### Learning the weights of top-down projections

So far, we have shown that supervised learning can be achieved in a dendritic EIANN that conforms to multiple biological constraints, including the *cellular diversity* constraint, the *signed synapse* constraint, and the *continuous signaling* constraint. However, in order to perform synaptic credit assignment comparable to backpropagation, the above results were obtained with top-down weights *B*^(*l*)^ set as symmetric with the bottom-up weights *W*^(*l* +1)^:

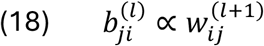

This violates the biological *non-symmetric synapse* constraint, which acknowledges that two different synapses in two different neurons do not share identical histories and may not undergo identical weight updates throughout training. One well-known solution proposed in previous work is to randomly initialize the top-down weights *B*^(*l*)^, and to simply fix them at their initial values throughout training^15^. This approach, called “feedback alignment,” has been shown to work surprisingly well in some supervised learning tasks, and works by encouraging the bottom-up weights *W*^(*l*+1)^ to gradually increase their symmetry with the top-down weights *B*^(*l*)^ over the course of learning. Related work has shown that the most important effect of this alignment is driven by symmetry in the signs of the *W*^(*l*+1)^ and *B*^(*l*)^ weights, even if their magnitudes are non-symmetric^87^. Interestingly, this symmetrical sign constraint is implicitly guaranteed by our *cellular diversity* and *signed synapse* constraints, which enforce that both the bottom-up and top-down connection weights are all excitatory (positive sign), resulting in partial alignment (45° angle) at initialization. However, unlike in this prior work, we found that fixing *B*^(*l*)^ caused alignment with *W*^(*l*+1)^ to progressively worsen over the course of training, rather than improve (Fig. 6b).

**Figure 6.**
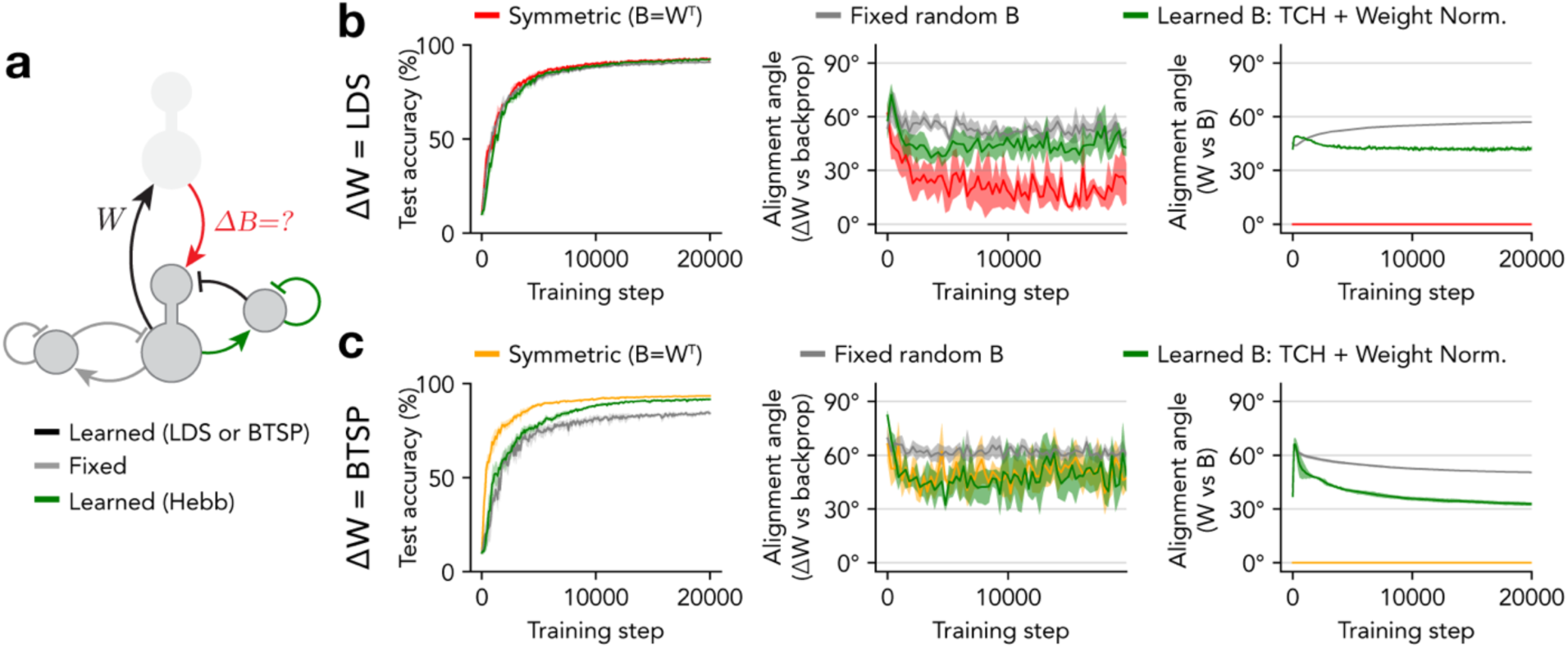
Independent learning of bottom-up and top-down synaptic weights. **a**, Diagram highlights that the bottom-up weights (*W*) and top-down weights (*B*) are distinct and may be learned independently. Full model includes projections to and from soma-targeting inhibitory neurons that are fixed and not learned (grey), projections onto dendrite-targeting inhibitory neurons that are learned with a Hebbian rule (green), and projections that are learned with dendritic state-dependent rules (black). **b**, Performance metrics for dendritic EIANN networks trained *with dendritic target propagation* with bottom-up weights (*W*) learned with the LDS rule and different strategies for learning the top-down weights (*B*): symmetric (red) sets *B* exactly symmetric to *W*, fixed random B (grey) fixes *B* at its random initialization, and learned B uses a temporally contrastive Hebbian (TCH) learning rule with weight normalization to update *B. Left*: Classification performance accuracy. *Middle*: Alignment angle between bottom-up excitatory (*W*) weight updates prescribed by each model and those prescribed by backpropagation. *Right*: Alignment angle between *W* and *B*. **c**, Same as **b** but for networks with bottom-up weights (*W*) learned with BTSP (symmetric model in yellow). **b-c**, Shading indicates standard deviation across five instances of each network.

The question of how to learn the top-down weights has also been considered by prior work, and most proposed solutions fall into two broad categories: 1) methods that learn *B*^(*l*)^ in a separate phase from *W*^(*l*+1)^, in which each neuron is driven by a statistically independent source of noise rather than sensory stimuli^16,88-90^, and 2) methods that simultaneously update both *B*^(*l*)^ and *W*^(*l*+1)^ by the same amount in each train step (sometimes referred to as the “Kolen-Pollack algorithm”)^14,16,26,91^. Neither of these approaches are fully biologically plausible, as real neurons share non-independent sources of noise (even during sleep periods), and transporting weight updates is a non-local operation. We therefore formulated an alternative solution that approximates the second approach while adhering more closely to biological constraints.

Since the top-down synapse 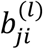 resides in the lower-layer neuron *j* (in layer *l*), it has local access to the activity 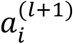 of the presynaptic upper-layer neuron *i* (in layer *l* + 1), but not to its dendritic state 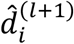, which is required to explicitly match the bottom-up update 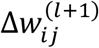 (Equation (9)). However, as we showed with the TCH rule (Equation 14 – 15), a neuron’s response to a dendritic “nudge” can be used to locally infer the magnitude of that nudge through a temporal comparison of its activity. Thus, we formulated a learning rule for the top-down weights *B*^(*l*)^ that leverages the same temporal comparison to align their updates with the dendritic signals that drive updates to the bottom-up weights *W*^(*l*+1)^:

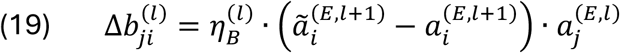

This is followed by a rescaling of the weights onto each postsynaptic neuron, as done in the Hebbian rule with weight normalization^62^ (Fig. 2c) (Equation (3)):

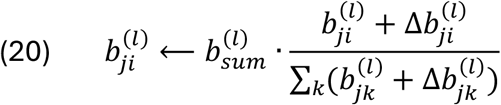

We found that learning top-down weights using this TCH rule with weight normalization resulted in improved symmetry of bottom-up weights *W*^(*l*+1)^ and top-down weights *B*^(*l*)^, compared to using fixed top-down weights (Fig. 6b). This improvement held across models using either the LDS or BTSP rule for learning the bottom-up weights (Fig. 6b,c). These results demonstrate that *dendritic target propagation* can be used to train multilayer deep neural networks while satisfying multiple biological constraints that have proved challenging to incorporate into ANN models, including the *non-symmetric synapse* constraint.

## Discussion

In this study, we imposed strict biological constraints on the architecture, information processing modes, and synaptic learning rules of multilayer artificial neural networks to better understand the biological mechanisms that coordinate changes in synaptic weights across multiple circuit layers during task learning. The design of our network and learning algorithm was informed by recent neuroscience experiments, which showed that top-down feedback signals from cortex provide target signals to instruct learning in the hippocampus^19^, and that this plasticity is mediated by dendritic calcium spikes initiated in the distal apical dendrites of excitatory pyramidal neurons^17,18^. Our *dendritic target propagation* algorithm implements separate soma and dendrite compartments in excitatory neurons, and computes plasticity-driving error signals locally in dendrites as the difference between top-down excitation and lateral inhibition from local dendrite-targeting interneurons. Our results demonstrate that using dendritic signaling to modulate somatic activity and synaptic plasticity is an effective biology-constrained mechanism to perform deep credit assignment, adjusting the right synapses in the right neurons to improve task performance, and relaying that information across multiple layers of neuronal circuitry.

Other recent theoretical work has proposed that biological neurons might perform credit assignment by multiplexing bottom-up sensory signals to neuronal soma compartments with top-down error signals to distal dendrites^26,27^. These models rely heavily on low-pass filtering bottom-up inputs to selectively pass “regular” spikes, and high-pass filtering top-down inputs to selectively pass bursts. They also consider dendritic activation to influence only the rate of bursts at the soma, but not the rate of regular spikes. It has not yet been demonstrated if those algorithms would be robust to the incorporation of the biological *signed synapse* constraint, or to relaxing the assumptions on synaptic filtering to match the more moderate and heterogenous dynamics observed experimentally^92^. However, that approach to deep credit assignment is complementary and not mutually exclusive to the one proposed in the current study, and it may be beneficial in the future to combine them, using both privileged processing of multiplexed spikes and bursts and gating of plasticity by local dendritic inhibition. Another line of research has focused on the advantages of nonlinear computations in basal and proximal apical dendrites for processing bottom-up inputs^93^, with theoretical work showing that single biological neurons with complex dendritic morphologies can classify input patterns comparably to entire deep networks^94,95^. That work is also complementary to ours, which focuses instead on the separate problem of processing top-down signals containing target information for synaptic credit assignment. Future dendritic EIANNs could be designed that both use *dendritic target propagation* for error computation, and contain more complex perisomatic dendrites for nonlinear processing of bottom-up sensory inputs.

An advantage of building computational models using components that directly correspond to biological counterparts is that the models can make precise predictions about specific circuit elements that can be tested experimentally. A key observation from our models is that using a learning rule to adjust both the synapses onto dendrite-targeting interneurons and their inhibitory connections onto the dendrites plays an important role in enabling the activation state of distal dendrites to approximate the local error gradient. While experiments have indicated that incoming and outgoing synapses to and from inhibitory cells can be plastic^96-99^, it has not yet been tested directly whether calcium spikes or dendritic voltage in pyramidal neurons modify dendritic inhibitory strengths, or whether the stimulus tuning of dendrite-targeting inhibitory cells adapts during learning. Interestingly, recent experiments have demonstrated that different dendritic compartments can express different synaptic learning rules^73,100-102^, which is consistent with the idea that learning in dendritic subcircuits might be specially tuned for optimal local error computation.

We note that, while the components of our model were heavily inspired by experimental observations and the microcircuit architectures of the hippocampus and neocortex, the layered network architecture of our model is generic, and is not meant to exactly mimic the architecture of any particular brain region. A natural extension of our work would be to more directly model the architectures of specific brain regions, for example by including recurrent lateral connections between excitatory neurons, as seen in cortical layer 2/3 or the CA3 region of the hippocampus. This could enable additional functionality, including selectivity for temporal sequences of inputs^103,104^, and the ability to retrieve associative memories from partial or degraded inputs^105,106^. Another possibility would be to more accurately model the top-down connectivity of the hippocampus, in which multiple circuit layers receive direct feedback projections from entorhinal cortex, rather than each layer receiving feedback only from adjacent layers^107,108^. Such models could help to inform the design of brain-region- and cell-type-specific circuit interventions for neurological disorders.

In this work, we demonstrate a biologically-plausible mechanism for the brain to approximate learning by gradient descent. However, to the extent that an algorithm precisely aligns with backpropagation, it will also suYer from the same drawbacks as backpropagation, including the well-known problem of “catastrophic forgetting,” where sequential training on multiple tasks causes degraded performance on previously learned tasks^109^. Interestingly, some previously proposed methods for reducing memory interference and improving task generalization involved imposing constraints on growth of synaptic weights, sparsity of population activity, or sparsity of gradients^110-113^. These regularization methods have direct analogies to many of the biological mechanisms implemented in our dendritic EIANNs, including weight normalization and regulation of activity and plasticity by feedback inhibition. In the future, it will be important to identify which aspects of biological learning allow the brain to mitigate memory interference and to learn efficiently from few training examples^114^.

In conclusion, our work leverages the computational power of biological neuronal microcircuits and neurons with compartmentalized signaling in dendrites to implement a deep learning algorithm that can solve nonlinear classification tasks in a manner fully compatible with principles of neuroscience. The recent discovery of behavioral timescale synaptic plasticity (BTSP) expanded the repertoire of known biological plasticity rules, and its unique features appear to offer advantages over previously characterized unsupervised learning rules, like spike timing-dependent plasticity (STDP), for target-directed learning^11^. However, the theoretical implications of BTSP for understanding and modeling biological learning have only begun to be explored^18,32,33^. Our results provide new insight into the role BTSP plays in enabling behavioral outcomes to drive synaptic changes distributed across multiple brain areas.

## Methods

### Network architecture and dynamics

Specification and training of ANN and EIANN models were performed in Python using a custom extension to PyTorch^115,116^. To train networks on the MNIST handwritten digit character dataset^117^, pixel intensities of image stimuli (28 × 28 pixels) were normalized between zero and one and encoded in the activities of 784 input neurons. Hidden layers (*l* = 1 and *l* = 2) each contained 500 excitatory neurons, 50 soma-targeting inhibitory neurons, and, when included, 50 dendrite-targeting inhibitory neurons. The output layer (*L* = 3) contained 10 excitatory neurons (one for each stimulus class) and 10 soma-targeting inhibitory neurons. For networks trained on the 2D spirals dataset, stimuli were encoded in the activities of 2 input neurons. Hidden layer *l* = 1 contained 128 excitatory neurons, 32 soma-targeting inhibitory neurons and, when included, 54 dendrite-targeting inhibitory neurons. Hidden layer *l* = 2 contained 32 excitatory neurons, 8 soma-targeting inhibitory neurons, and when included, 13 dendrite-targeting inhibitory neurons. The output layer (*L* = 3) contained 4 excitatory neurons (one for each stimulus class) and 4 soma-targeting inhibitory neurons.

During training, stimuli were presented to a network one at a time (batch size of 1) to mimic serial sampling of the sensory environment by biological organisms. For networks with recurrent (lateral) connections between separate excitatory and inhibitory cells (EIANNs), stimulus inputs were held constant for 15 time steps during a sensory phase while the somatic states and output firing rates of the neurons in all layers were equilibrated (Supplementary Fig. S1). Excitatory neuron somatic states (*S*^(*E,l*)^) were updated in each time step as:

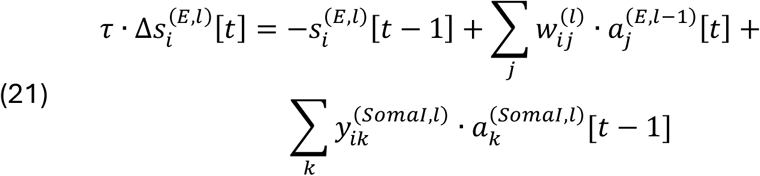

where *τ* = 3 is a time constant. Soma-targeting inhibitory neuron somatic states were updated as:

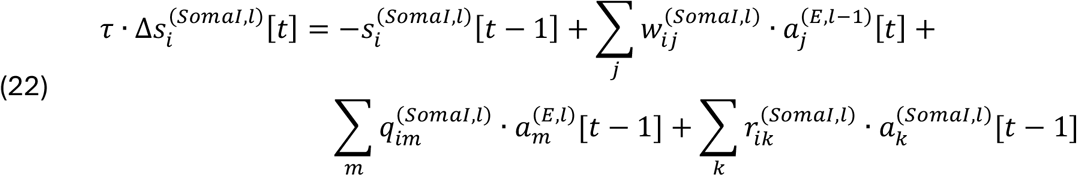

Dendrite-targeting inhibitory neuron somatic states were updated as:

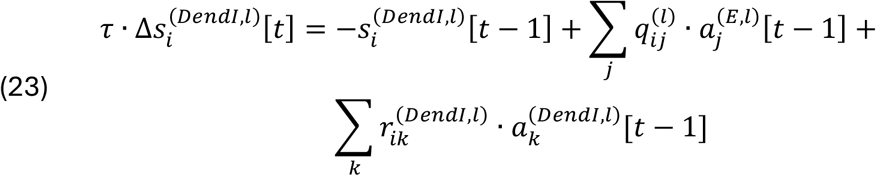

The weights of self-connections (*i* = *k*) between recurrently connected interneurons 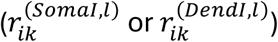 were set to zero. Finally, the ReLU activation function was used to determine the activities of neurons from their somatic states:

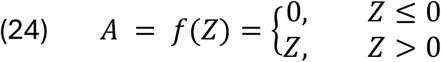

At the end of the sensory phase, output layer dendritic states 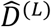 and activities 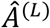 were updated by a supervisory “nudge” according to Equations (6) – (8). In hidden layers, the “nudged” dendritic states were then computed as follows:

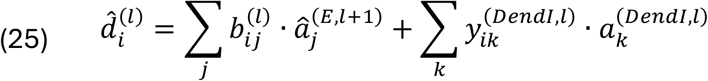

Dendritic states 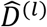 were then bounded between -1 and 1 and held constant during the supervised phase for an additional 15 time steps while the somatic states 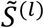 and firing rates *Ã* ^(*l*)^ of all neurons re-equilibrated according to Equations (21) – (24) and (8).

### Learning rules

When backpropagation was used to learn the bottom-up excitatory weights *W*^(*E,l*)^ (Fig. 2a,b and Fig. 4), a global loss function was defined as the mean squared error of the output neurons relative to a one-hot target vector:

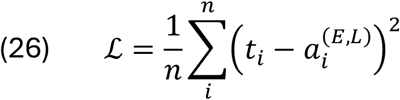

Then, each synaptic weight was updated proportionally to the gradient of loss with respect to the weight:

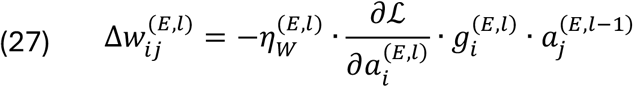

In networks with EIANN recurrent architectures, the loss gradient was backpropagated through 3 equilibration time steps. When the incoming and outgoing projections of soma-targeting interneurons were learned by backpropagation (Fig. 2b, and Supplementary Fig. S1 – S3), the bottom-up and lateral weights (*W*^(*somal,l*)^, *Q*^(*somal,l*)^, and *R*^(*somal,l*)^) were updated using the same method.

In Fig. 2c, the Hebbian learning rule with weight normalization described in Equations (1) – (3) was used to update the bottom-up (*W*^(*E,l*)^) and lateral (*Y*^(*somal,l*)^) weights onto excitatory neurons and the bottom-up (*W*^(*somal,l*)^) and lateral (*Q*^(*somal,l*)^ and *R*^(*somal,l*)^) weights onto soma-targeting inhibitory neurons in the hidden layers. For that network, the bottom-up excitatory weights *W*^(*E,L*)^ onto output layer excitatory neurons were updated using the error-dependent rule described in Equations (6) – (9). The bottom-up (*W*^(*somal,l*)^) and lateral (*Q*^(*somal,l*)^ and *R*^(*somal,l*)^) weights onto soma-targeting inhibitory neurons in the output layer were updated according to Equations (1) – (3,) after activities *Ã* ^(*E,L*)^ and *Ã* (^*somal,l*^) had equilibrated during the supervised phase for an additional 15 time steps after the “nudge” was applied to the excitatory neurons.

When the BCM rule was used to update the bottom-up excitatory weights *W*^(*El*)^ using Equation (16) (Fig. 5b), sliding plasticity thresholds *θ*^(*l*)^ were updated after each stimulus presentation as:

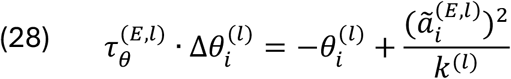

where 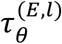 is a time constant, *k*^(*l*)^ is a constant scale factor, and 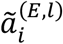 is the activity of postsynaptic neuron *i* after supervisory inputs have been applied and the network has re-equilibrated.

The BTSP rule defined in Equation (17) refers to presynaptic eligibility traces *ET*^(*l* − 1)^ and postsynaptic instructive signals *IS*^(*l*)^. When both presynaptic activity 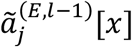 and postsynaptic dendritic state 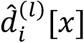 were greater than zero in response to the current stimulus *x*, then 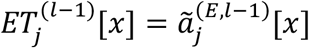 (bounded to be <= 1), 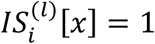, and the weight 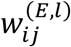 was updated according to Equation (17). The sigmoidal function *σ*(*z*) referenced in Equation (17) is scaled and offset to meet the edge constraints *σ*(0) = 0 and *σ*(1) = 1:

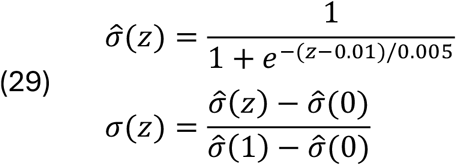

The BTSP rule also prescribes weight updates for inputs active at long time delays from a dendritic calcium spike^18^. Here we modeled this aspect in the following way: when a presynaptic unit *j* was active in response to the previous stimulus *x* − 1, then the eligibility trace 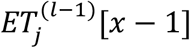 was decayed by a factor of *λ*^(*l*)^ by the time of stimulus *x*, resulting in 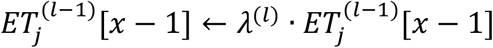 and an additional weight update according to:

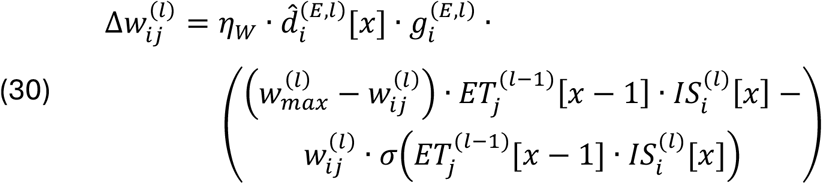

When a postsynaptic unit’s dendritic state 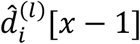 in response to the previous stimulus *x* − 1 was greater than zero, then the instructive signal 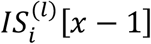 was decayed by a factor of *λ*^(*l*)^ by the time of stimulus *x*, resulting in 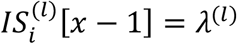 and an additional weight update according to:

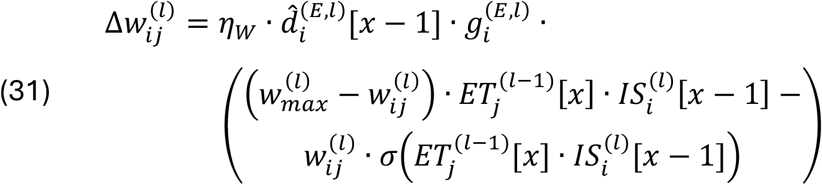

Finally, while the experimentally-observed BTSP learning rule is defined when calcium spikes are triggered by dendritic depolarization above rest (positive dendritic state), it is not known how weights may be updated when dendrites are hyperpolarized below rest (negative dendritic state). In this case when 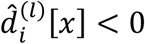, weights were updated using the LDS rule described in Equation (9).

### Network training and hyperparameter optimization

The weights of each synaptic projection between cell types and layers were initialized from uniform distributions, respecting the *signed synapse* constraint, with projection-specific scales selected during hyperparameter optimization (discussed below; see Supplementary Table S3 – S6). During training, five instances of each network were separately initialized using different random seeds, and separately trained on the same stimulus samples, but presented in a different shuffled order. For the MNIST handwritten digit classification task, each network instance was trained by serially presenting either 20,000 or 50,000 images (Supplementary Table S1) without repeat (less than a single epoch of the dataset), with one weight update applied after each image. Final task accuracy was evaluated by presenting a separate test set of 10,000 images not used during training, and quantified as the percent of test samples where the most active (excitatory) output neuron corresponded to the correct stimulus class. During hyperparameter optimization, task accuracy was evaluated by presenting a separate validation set of 10,000 images not used during either training or final testing, and averaging the validation accuracy across the five network instances. For the 2D spirals task, networks were trained for either 1 or 10 epochs on a train set of 5,600 samples (Supplementary Table S2). Accuracy was evaluated with distinct train and validation sets of 1,200 samples each.

Each network was specified by a set of hyperparameters that included projection-specific scales for weight initialization or weight normalization, projection-specific learning rates, and parameters internal to each synaptic learning rule (Supplementary Table S3 – S6). For each network, these hyperparameters were searched within specified bounds using custom Python code implementing an iterative population-based search based on the basin-hopping algorithm^118,119^ that evaluated 30,000 variants of each network. Optimization aimed to minimize the following objectives, each computed as an average across the five instances of each network variant: 1) MSE loss (Equation (26)); 2) validation accuracy error (100% - validation accuracy); 3) sensory-phase dendritic state, averaged across cells; and 4) residual oscillations in activity after sensory-phase equilibration, averaged across cells (Supplementary Figure S1). The resulting hyperparameters for each network are shown in Supplementary Table S3 – S6. For an excitatory projection with *n* presynaptic neurons and weights *W*^(*l*)^, a hyperparameter 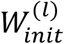 was used to initialize weights from a uniform distribution between zero and 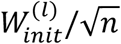. When top-down weights *B*^(*l*)^ were set as symmetric to bottom-up weights *W*^(*l*+1)^, a hyperparameter 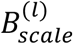 was used to rescale the top-down weights as 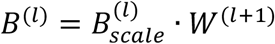.

### Analysis of neuronal stimulus selectivity

The stimulus selectivity of a neuron (Fig. 2 – 4 and Supplementary Fig. S2 and S4) was quantified as:

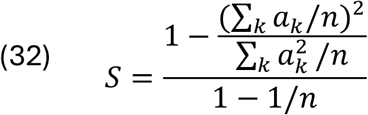

where *a*_*k*_ is the activity in response to stimulus *k*, and *n* is the number of stimuli in a dataset^120^. A value of zero corresponds to uniform responses to all stimuli, and a value of one indicates that a unit responds maximally to a single stimulus and minimally to all other stimuli.

For networks trained to classify handwritten digits, we also analyzed the stimulus preferences of individual neurons by visualizing their receptive fields in image space using an algorithm adapted from previous work^50^ (Fig. 2 and 4 and Supplementary Fig. S3 – S5). We selected 1000 random images from the MNIST test dataset and presented them to trained networks with frozen parameters. For each unit in the network, we defined a loss function equal to the negative of the unit’s activity, and computed the gradient of this loss function with respect to each pixel in each presented image. These gradients were then averaged over the 1000 images and normalized, resulting in a visualization representing the average positive or negative influence of each pixel on the unit’s activity. We quantified the spatial structure of these two-dimensional receptive fields (Fig. 2 and 4 and Supplementary Fig. S4) using Moran’s I measure of spatial autocorrelation:

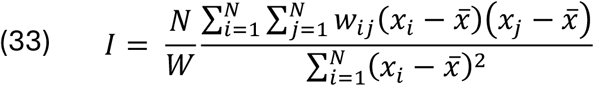

where *N* is the number of pixels, *x*_*i*_ and *x*_*j*_ are the intensities of a pair of pixels *i* and *j*, 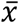 is the mean pixel intensity, *w*_*ij*_ is a spatial *weight* between a pair of pixels, and *W* is the sum of all weights *w*_*ij*_. The weights were set to one for neighboring pixels and zero otherwise, resulting in *I* values close to zero for random noise images and approaching one for images with strong correlations across pixels within spatial domains.

## Acknowledgements

This work was supported by funding from Rutgers Biomedical and Health Sciences (A.P., A.D.M.), NIMH grant R01MH121979 (A.P., A.R.G., A.D.M.), NIMH grant R01MH135576 (A.R.G., A.D.M.), EMBO postdoctoral fellowship ALTF 473-2022 (A.R.G.), and Rutgers Aresty Research Center (Y.C.). Large-scale compute resources were provided by Rutgers OARC (Amarel) and NSF-supported allotments from TACC (Frontera) and ACCESS (Bridges-2). We appreciated the opportunity to discuss and work out early concepts and code with students Tyler Rivera, Sandhya Senthilkumar, Barbara Gruszka, Maya Salameh, Michael Finch, Anshul Voleti, and Michael Cheng.

## Author contributions

Conceptualization: A.R.G., A.P., A.D.M.; supervision: A.D.M.; computational modeling: A.R.G., A.P., Y.C., A.D.M.; data analysis: A.R.G., Y.C., A.D.M.; writing—original draft: A.R.G., A.D.M.; writing—review & editing: A.R.G., A.D.M.

## Supplementary Figures

**Supplementary Figure S1.**
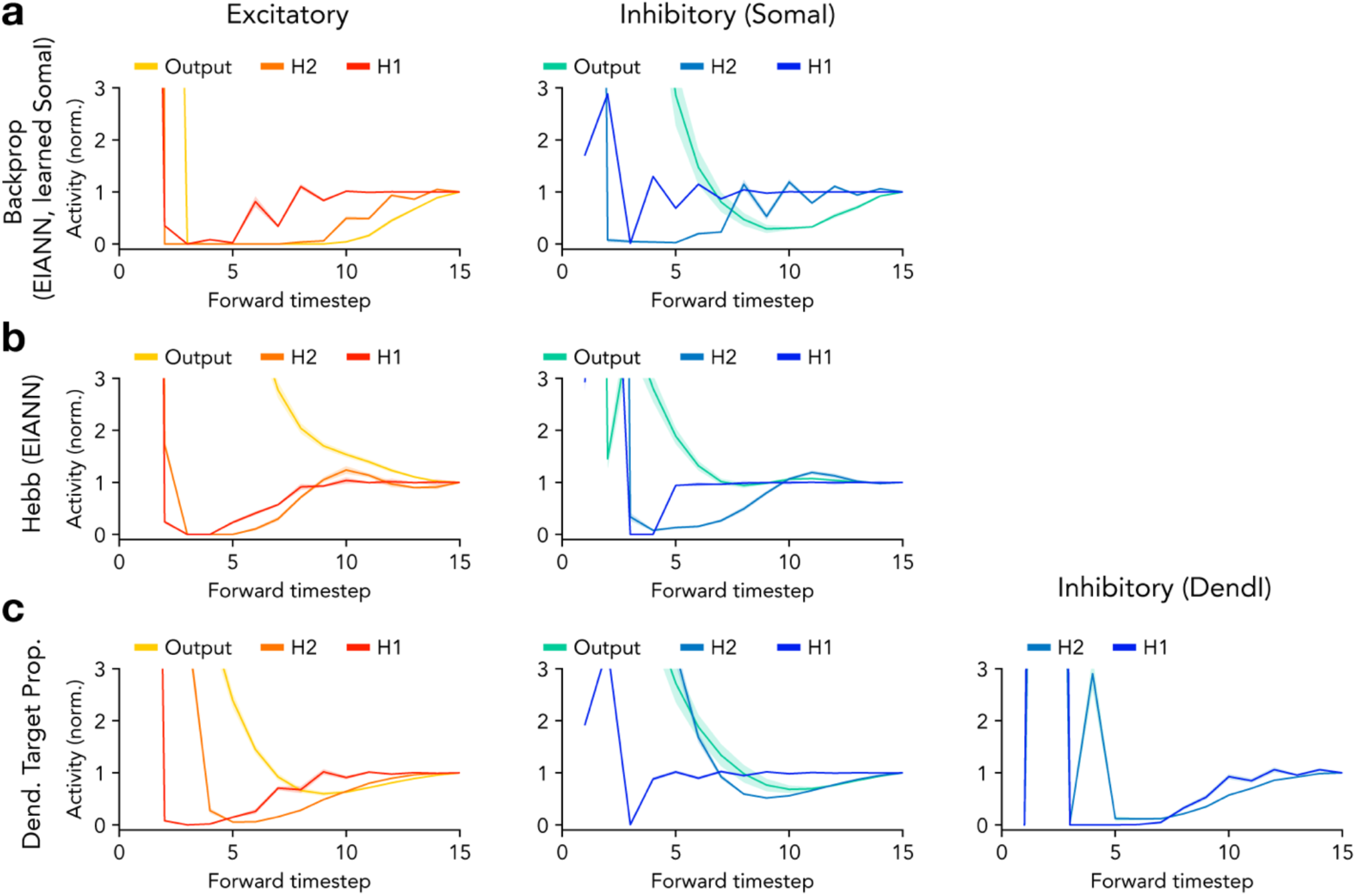
Equilibration dynamics of neuronal activity in recurrent EIANNs. Graphs show the dynamics of neuronal activity across 15 time steps of equilibration. To compare neuronal subpopulations, the average activity of each population is normalized such that its final value at the end of equilibration is one. Large onset transients are truncated for display purposes. *Left*: excitatory neurons; *middle*: soma-targeting inhibitory neurons (SomaI); *right*: dendrite-targeting inhibitory neurons (DendI). **a**, EIANN trained with backpropagation. **b**, EIANN trained with the normalized Hebbian learning rule. **c**, Dendritic EIANN trained with *dendritic target propagation*. **a-c**, Shading indicates standard deviation across five instances of each network.

**Supplementary Figure S2.**
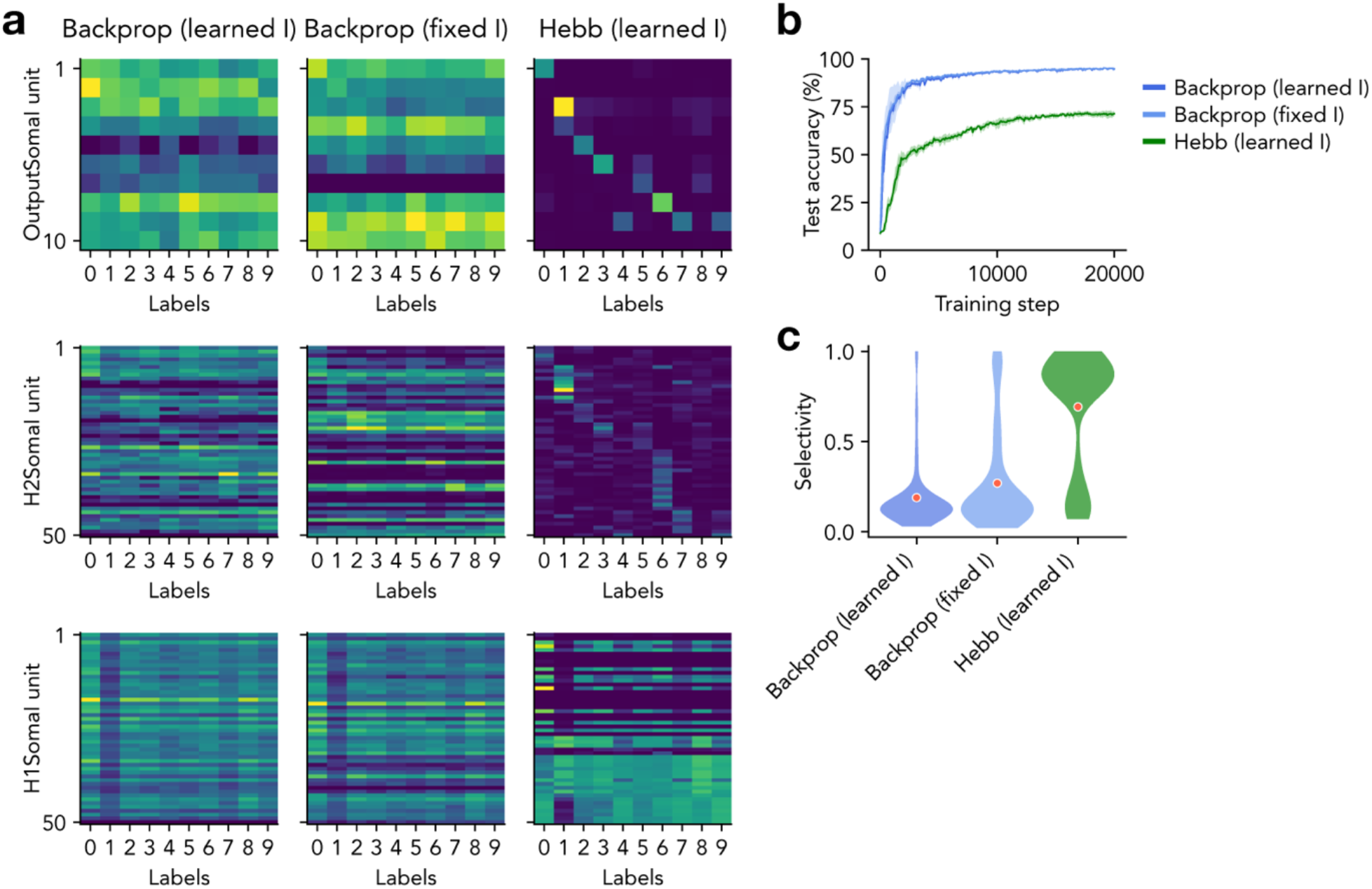
Plasticity of somatic inhibition is not required for image classification. **a**, Average neuronal population activity in response to each class of handwritten digits. *Left*: Shown are soma-targeting inhibitory neurons (SomaI) from the Output layer (10 units, top), layer H2 (50 units, middle), and layer H1 (50 units, bottom) from a recurrent EIANN trained with backpropagation with learned connections to and from SomaI units. *Middle*: Same as *left* for a recurrent EIANN trained with backpropagation with connections to and from SomaI units that are fixed at initialization and not learned. *Right*: same as *left* for a recurrent EIANN trained with a normalized Hebbian learning rule. **b**, Classification performance accuracy. Shading indicates standard deviation across five instances of each network. **c**, Selectivity of SomaI units (in all layers) over stimulus classes.

**Supplementary Figure S3.**
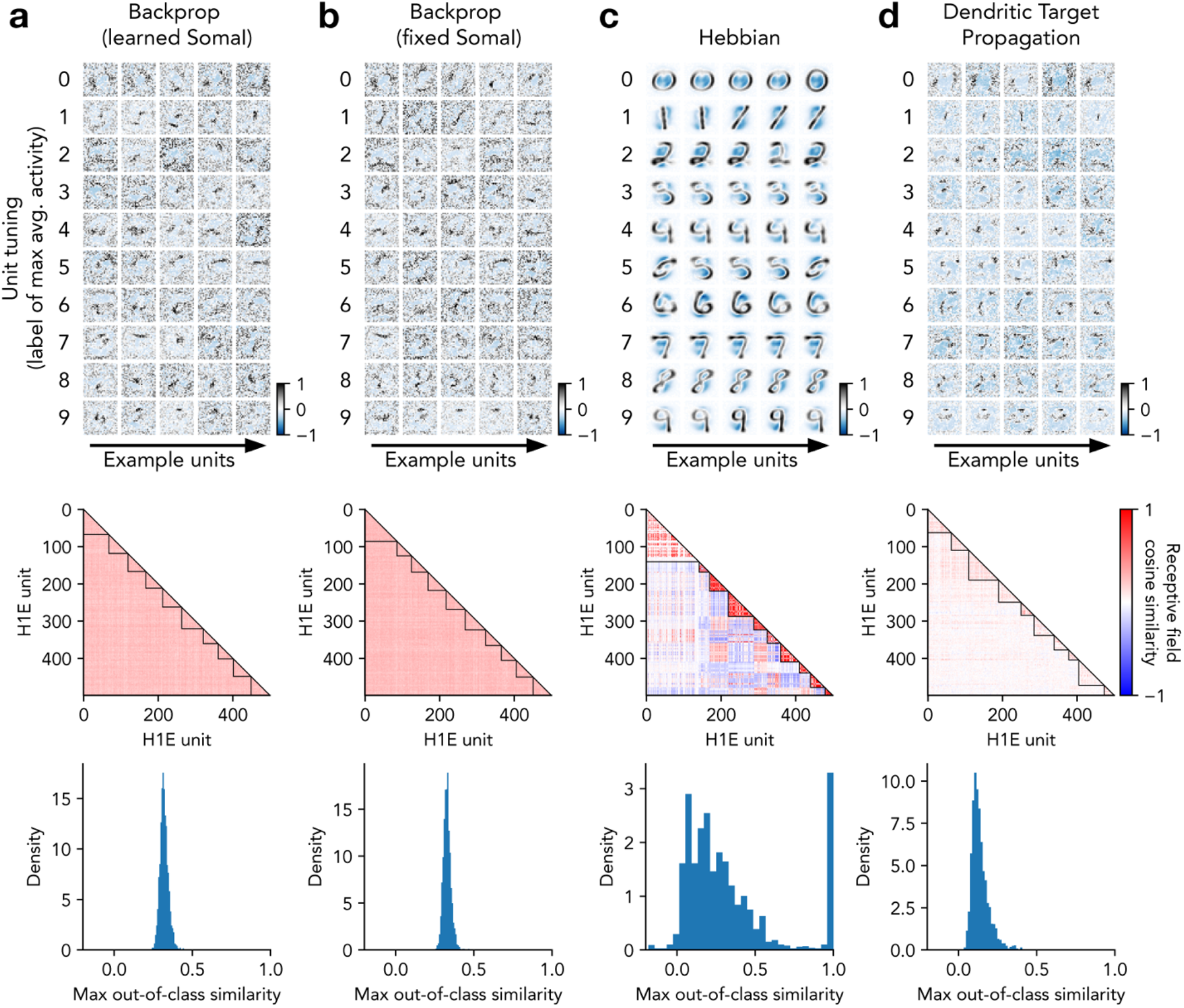
Analysis of hidden layer neuronal receptive fields in EIANNs. **a**, Receptive fields are analyzed from H1E units from a recurrent EIANN trained with backpropagation with learned connections to and from SomaI units. *Top*: Example receptive fields. Row labels indicate the class that produces the largest activity (averaged across samples) for each unit. *Middle*: Heatmaps depict the cosine similarity of receptive fields between pairs of H1E units. Units are sorted by their preferred stimulus class, and black lines demarcate groups of units that share a class preference. *Bottom*: Histograms show, for each unit, the maximum cosine similarity compared to other units that do not share the same stimulus class preference. **b**, Same as **a** for a recurrent EIANN trained with backpropagation with connections to and from SomaI units that are fixed at initialization and not learned. **c**, same as **a** for a recurrent EIANN trained with a normalized Hebbian learning rule. **d**, same as **a** for a dendritic EIANN trained with *dendritic target propagation*.

**Supplementary Figure S4.**
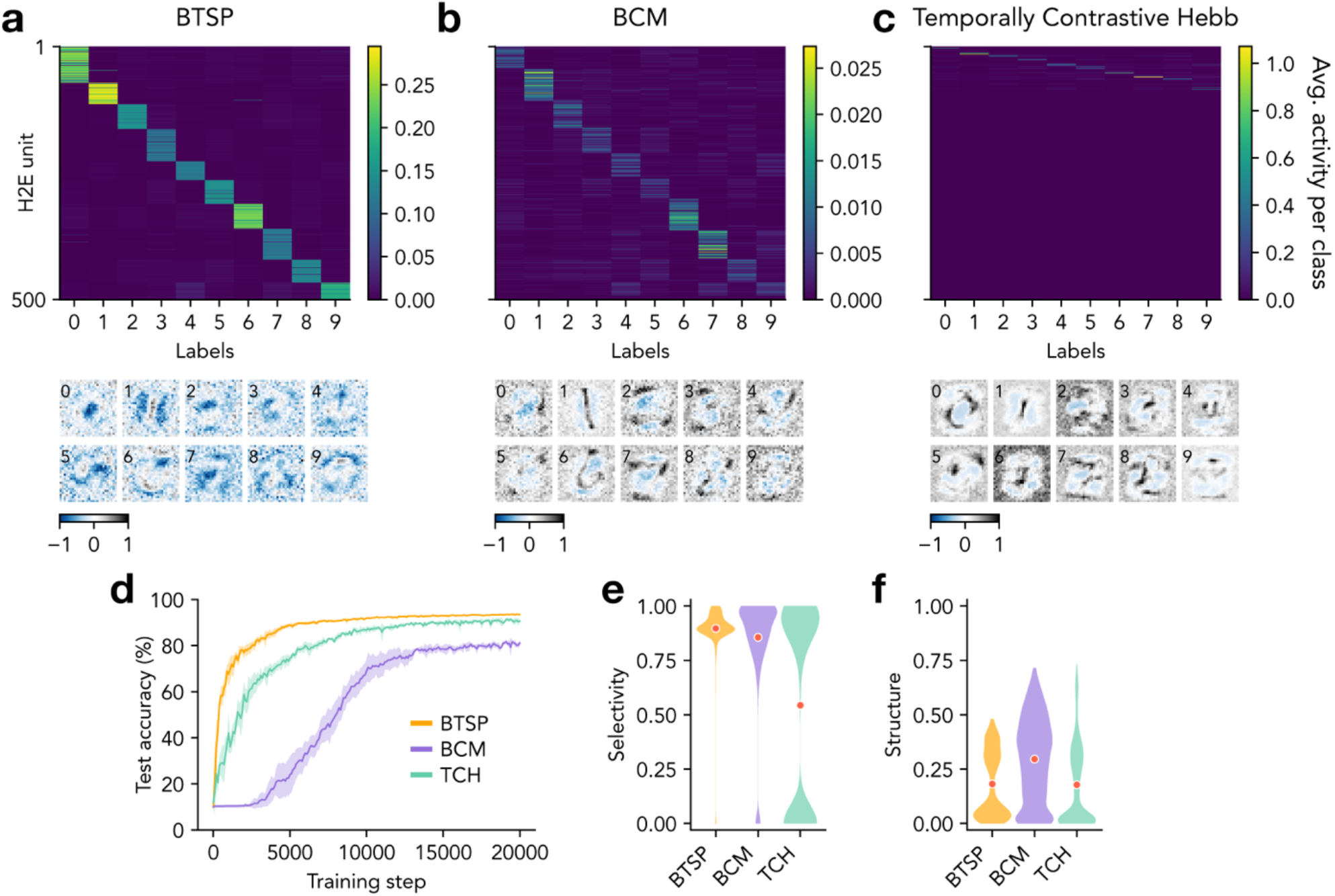
Excitatory neuron selectivity in dendritic EIANNs trained with biological learning rules. **a**, *Top*: Average neuronal population activity in response to each class of handwritten digits. Shown are 500 excitatory (E) units in the second hidden layer (H2) from a dendritic EIANN trained with *dendritic target propagation* and using the BTSP learning rule to learn bottom-up excitatory weights. *Bottom:* Example receptive fields selected from H2E units. Numbers in the top left corners indicate the label of the class that produces the largest activity (averaged across samples) for each unit. **b**, Same as **a** for a dendritic EIANN using the BCM rule to learn bottom-up excitatory weights. **c**, Same as **a** for a dendritic EIANN using the TCH rule to learn bottom-up excitatory weights. **d**, Classification performance accuracy. Shading indicates standard deviation across five instances of each network. **e**, Selectivity of E units (in both hidden layers) over stimulus classes. **f**, Spatial structure of the receptive fields of E units (in both hidden layers), as measured by spatial autocorrelation (Moran’s I).

**Supplementary Figure S5.**
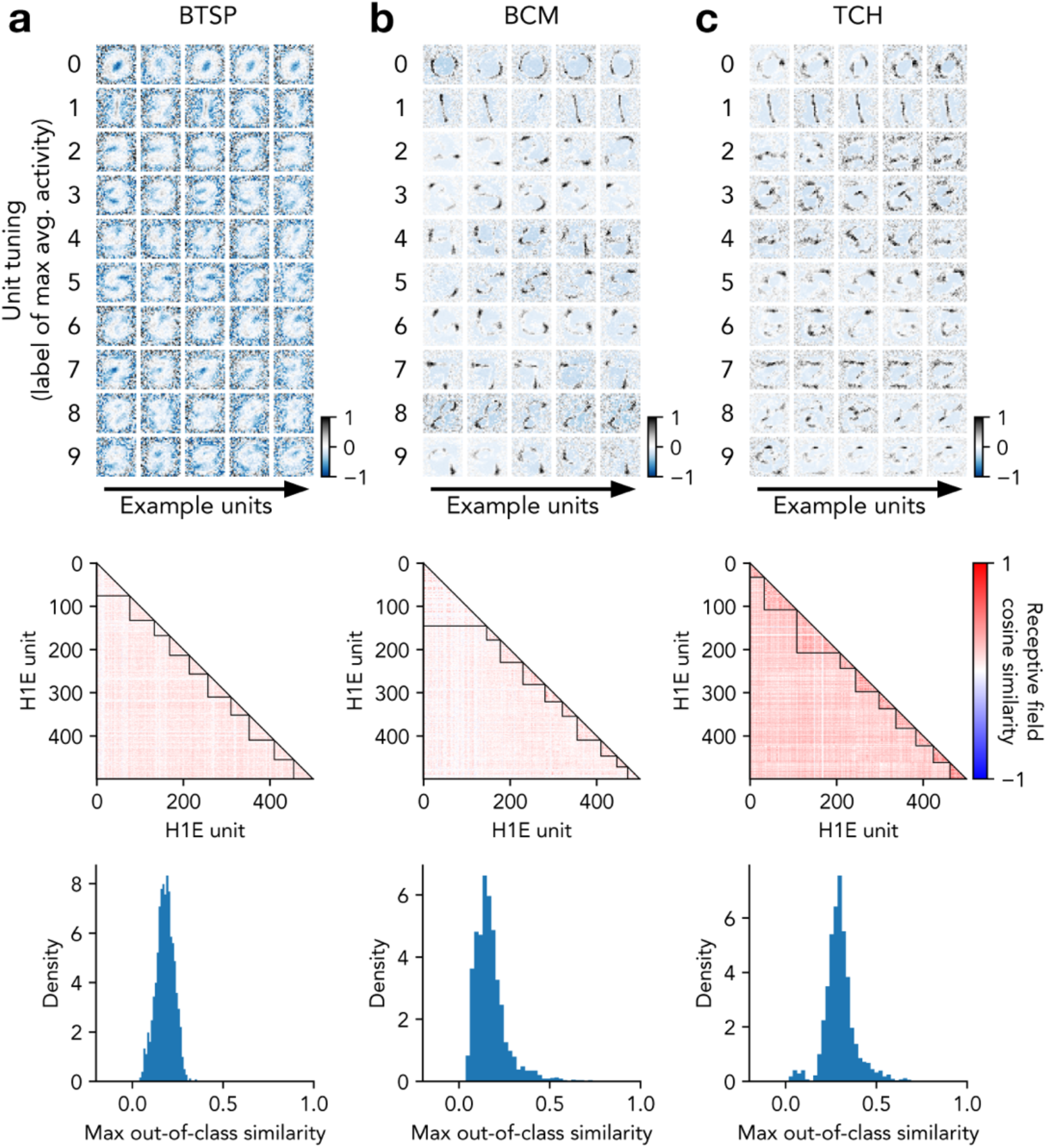
Analysis of hidden layer neuronal receptive fields in dendritic EIANNs trained with biological learning rules. **a**, Receptive fields are analyzed from H1E units from a dendritic EIANN trained with *dendritic target propagation* and using the BTSP learning rule to learn bottom-up excitatory weights. *Top*: Example receptive fields. Row labels indicate the class that produces the largest activity (averaged across samples) for each unit. *Middle*: Heatmaps depict the cosine similarity of receptive fields between pairs of H1E units. Units are sorted by their preferred stimulus class, and black lines demarcate groups of units that share a class preference. *Bottom*: Histograms show, for each unit, the maximum cosine similarity compared to other units that do not share the same stimulus class preference. **b**, Same as **a** for a dendritic EIANN using the BCM rule to learn bottom-up excitatory weights. **c**, Same as **a** for a dendritic EIANN using the TCH rule to learn bottom-up excitatory weights.

**Supplementary Table S1.** Handwritten digit classification performance accuracy.

|  | Architecture | Hidden Layers | Algorithm | W Learning Rule | B Learning Rule | Bias | MNIST Accuracy<br>(20k samples) | MNIST Accuracy<br>(50k samples) |
| --- | --- | --- | --- | --- | --- | --- | --- | --- |
| Feedforward ANN | ANN | 2 | Backprop | Gradient descent | Symmetric ($B = W^T$) | Zero | 95.48 ± 0.20 | 95.48 ± 0.20 |
| Feedforward ANN<br>(fixed hidden) | ANN | 2 | Backprop | Gradient descent | Symmetric ($B = W^T$) | Zero | 87.73 ± 0.08 | 89.68 ± 0.24 |
| Feedforward ANN<br>(no hidden) | ANN | 0 | Backprop | Gradient descent | Symmetric ($B = W^T$) | Zero | 89.43 ± 0.08 | 90.56 ± 0.05 |
| Backprop<br>(fixed Soma) | EIANN | 2 | Backprop | Gradient descent | Symmetric ($B = W^T$) | Zero | 94.70 ± 0.38 | 96.38 ± 0.16 |
| Backprop<br>(learned Soma) | EIANN | 2 | Backprop | Gradient descent | Symmetric ($B = W^T$) | Zero | 94.76 ± 0.27 | 96.53 ± 0.10 |
| Backprop<br>(no Soma) | EIANN | 2 | Backprop | Gradient descent | Symmetric ($B = W^T$) | Zero | 60.65 ± 1.72 | 64.01 ± 1.03 |
| Hebbian | EIANN | 2 | Unsupervised<br>(no propagation) | Hebb<br>+ Weight Norm. | Symmetric ($B = W^T$) | Zero | 71.35 ± 0.64 | 73.24 ± 0.60 |
| Fixed Dendl<br>(random) | Dend EIANN | 2 | Dend Target Prop | LDS | Symmetric ($B = W^T$) | Zero | 86.80 ± 0.50 | 87.36 ± 0.59 |
| Learned Dendl<br>(local backprop) | Dend EIANN | 2 | Dend Target Prop<br>+ Backprop (Dendl) | LDS | Symmetric ($B = W^T$) | Zero | 94.08 ± 0.11 | 95.04 ± 0.20 |
| Learned Dendl<br>(Hebb) | Dend EIANN | 2 | Dend Target Prop | LDS | Symmetric ($B = W^T$) | Zero | 92.62 ± 0.10 | 94.29 ± 0.13 |
| Temporally<br>Contrastive Hebb | Dend EIANN | 2 | Dend Target Prop | TCH | Symmetric ($B = W^T$) | Zero | 90.55 ± 0.30 | 90.89 ± 0.68 |
| BCM | Dend EIANN | 2 | Dend Target Prop | BCM | Symmetric ($B = W^T$) | Zero | 81.10 ± 0.52 | 68.92 ± 0.89 |
| BTSP | Dend EIANN | 2 | Dend Target Prop | BTSP | Symmetric ($B = W^T$) | Zero | 93.43 ± 0.10 | 94.08 ± 0.13 |
| LDS,<br>fixed top-down | Dend EIANN | 2 | Dend Target Prop | LDS | Fixed random | Zero | 90.93 ± 0.33 | 92.86 ± 0.19 |
| LDS,<br>learned top-down | Dend EIANN | 2 | Dend Target Prop | LDS | TCH + Weight Norm. | Zero | 92.47 ± 0.13 | 93.96 ± 0.12 |
| BTSP,<br>fixed top-down | Dend EIANN | 2 | Dend Target Prop | BTSP | Fixed random | Zero | 84.24 ± 0.47 | 83.32 ± 0.47 |
| BTSP,<br>learned top-down | Dend EIANN | 2 | Dend Target Prop | BTSP | TCH + Weight Norm. | Zero | 91.65 ± 0.17 | 93.50 ± 0.14 |

**Supplementary Table S2.** Two-dimensional spiral pattern classification performance accuracy.

|  | Architecture | Hidden Layers | Algorithm | W Learning Rule | B Learning Rule | Bias | Spiral Accuracy<br>(1 epoch) | Spiral Accuracy<br>(10 epochs) |
| --- | --- | --- | --- | --- | --- | --- | --- | --- |
| Feedforward ANN<br>(no hidden) | ANN | 0 | Backprop | Gradient Descent | Symmetric ($B = W^T$) | Learned | 55.35 ± 2.64 | 51.38 ± 2.17 |
| Feedforward ANN<br>(learned bias) | ANN | 2 | Backprop | Gradient Descent | Symmetric ($B = W^T$) | Learned | 95.77 ± 0.41 | 97.82 ± 0.10 |
| Feedforward ANN<br>(no bias) | ANN | 2 | Backprop | Gradient Descent | Symmetric ($B = W^T$) | Zero | 48.95 ± 0.07 | 48.58 ± 0.16 |
| Backprop (EIANN) | EIANN | 2 | Backprop | Gradient Descent | Symmetric ($B = W^T$) | Learned | 95.87 ± 0.27 | 97.08 ± 0.35 |
| Dendritic<br>Target Propagation | Dend EIANN | 2 | Dend Target Prop | LDS | Symmetric ($B = W^T$) | Learned | 91.87 ± 0.74 | 90.13 ± 1.04 |

**Supplementary Table S3.** Hyperparameters for networks trained on handwritten digit classification (Part 1).

| Param name | Feedforward ANN | Feedforward ANN<br>(fixed hidden) | Feedforward ANN<br>(no hidden) | Backprop<br>(fixed Soma) | Backprop<br>(learned Soma) | Backprop<br>(no Soma) | Hebbian |
| --- | --- | --- | --- | --- | --- | --- | --- |
| $W_{init}$ (H1) | 2.9573 | 5.0120 | - | 2.4169 | 2.3418 | 0.0313 | - |
| $\eta$, W (H1) | 0.1652 | - | - | 0.3981 | 0.3913 | 0.1904 | 0.0055 |
| $W_{init}$ (H2) | 0.3069 | 0.3272 | - | 0.5428 | 0.7159 | 1.4197 | - |
| $\eta$, W (H2) | 0.1652 | - | - | 0.3981 | 0.3913 | 0.1904 | 0.0055 |
| $W_{init}$ (Output) | 2.0936 | 0.0203 | 0.0265 | 3.2817 | 1.0173 | 3.9650 | 1.1091 |
| $\eta$, W (Output) | 0.0217 | 0.0582 | 0.0041 | 0.0708 | 0.3265 | 0.0110 | 0.0193 |
| $Y_{init}$ (Soma, H1) | - | - | - | 1.1159 | 1.5332 | - | - |
| $W_{init}$ (Soma, H1) | - | - | - | 3.1371 | 2.0286 | - | - |
| $Q_{init}$ (Soma, H1) | - | - | - | 0.4836 | 0.3540 | - | - |
| $R_{init}$ (Soma, H1) | - | - | - | 1.2959 | 1.2781 | - | - |
| $Y_{init}$ (Soma, H2) | - | - | - | 0.4713 | 0.9795 | - | - |
| $W_{init}$ (Soma, H2) | - | - | - | 1.4413 | 1.0560 | - | - |
| $Q_{init}$ (Soma, H2) | - | - | - | 1.5897 | 0.1488 | - | - |
| $R_{init}$ (Soma, H2) | - | - | - | 2.4085 | 1.4983 | - | - |
| $Y_{init}$ (Soma, Output) | - | - | - | 1.5076 | 0.9429 | - | - |
| $W_{init}$ (Soma, Output) | - | - | - | 2.9499 | 0.8592 | - | - |
| $Q_{init}$ (Soma, Output) | - | - | - | 0.5984 | 0.2251 | - | - |
| $R_{init}$ (Soma, Output) | - | - | - | 1.5326 | 0.8824 | - | - |
| $\eta$, Y (Soma, H1) | - | - | - | - | 0.0016 | - | 0.0388 |
| $\eta$, W (Soma, H1) | - | - | - | - | 0.0851 | - | 0.0007 |
| $\eta$, Q (Soma, H1) | - | - | - | - | 0.0851 | - | 0.0007 |
| $\eta$, R (Soma, H1) | - | - | - | - | 0.0029 | - | 0.0523 |
| $\eta$, Y (Soma, H2) | - | - | - | - | 0.0016 | - | 0.0388 |
| $\eta$, W (Soma, H2) | - | - | - | - | 0.0851 | - | 0.0007 |
| $\eta$, Q (Soma, H2) | - | - | - | - | 0.0851 | - | 0.0007 |
| $\eta$, R (Soma, H2) | - | - | - | - | 0.0029 | - | 0.0523 |
| $\eta$, Y (Soma, Output) | - | - | - | - | 0.0016 | - | 0.0388 |
| $\eta$, W (Soma, Output) | - | - | - | - | 0.0851 | - | 0.0007 |
| $\eta$, Q (Soma, Output) | - | - | - | - | 0.0851 | - | 0.0007 |
| $\eta$, R (Soma, Output) | - | - | - | - | 0.0029 | - | 0.0523 |
| $W_{sum}$ (H1) | - | - | - | - | - | - | 9.6360 |
| $Y_{sum}$ (Soma, H1) | - | - | - | - | - | - | 5.6391 |
| $W_{sum}$ (Soma, H1) | - | - | - | - | - | - | 13.5545 |
| $Q_{sum}$ (Soma, H1) | - | - | - | - | - | - | 14.2162 |
| $R_{sum}$ (Soma, H1) | - | - | - | - | - | - | 5.7558 |
| $W_{sum}$ (H2) | - | - | - | - | - | - | 6.9501 |
| $Y_{sum}$ (Soma, H2) | - | - | - | - | - | - | 2.4875 |
| $W_{sum}$ (Soma, H2) | - | - | - | - | - | - | 6.1176 |
| $Q_{sum}$ (Soma, H2) | - | - | - | - | - | - | 7.3201 |
| $R_{sum}$ (Soma, H2) | - | - | - | - | - | - | 2.6716 |
| $Y_{sum}$ (Soma, Output) | - | - | - | - | - | - | 0.1380 |
| $W_{sum}$ (Soma, Output) | - | - | - | - | - | - | 5.5863 |
| $Q_{sum}$ (Soma, Output) | - | - | - | - | - | - | 1.0857 |
| $R_{sum}$ (Soma, Output) | - | - | - | - | - | - | 2.8848 |

**Supplementary Table S4.**
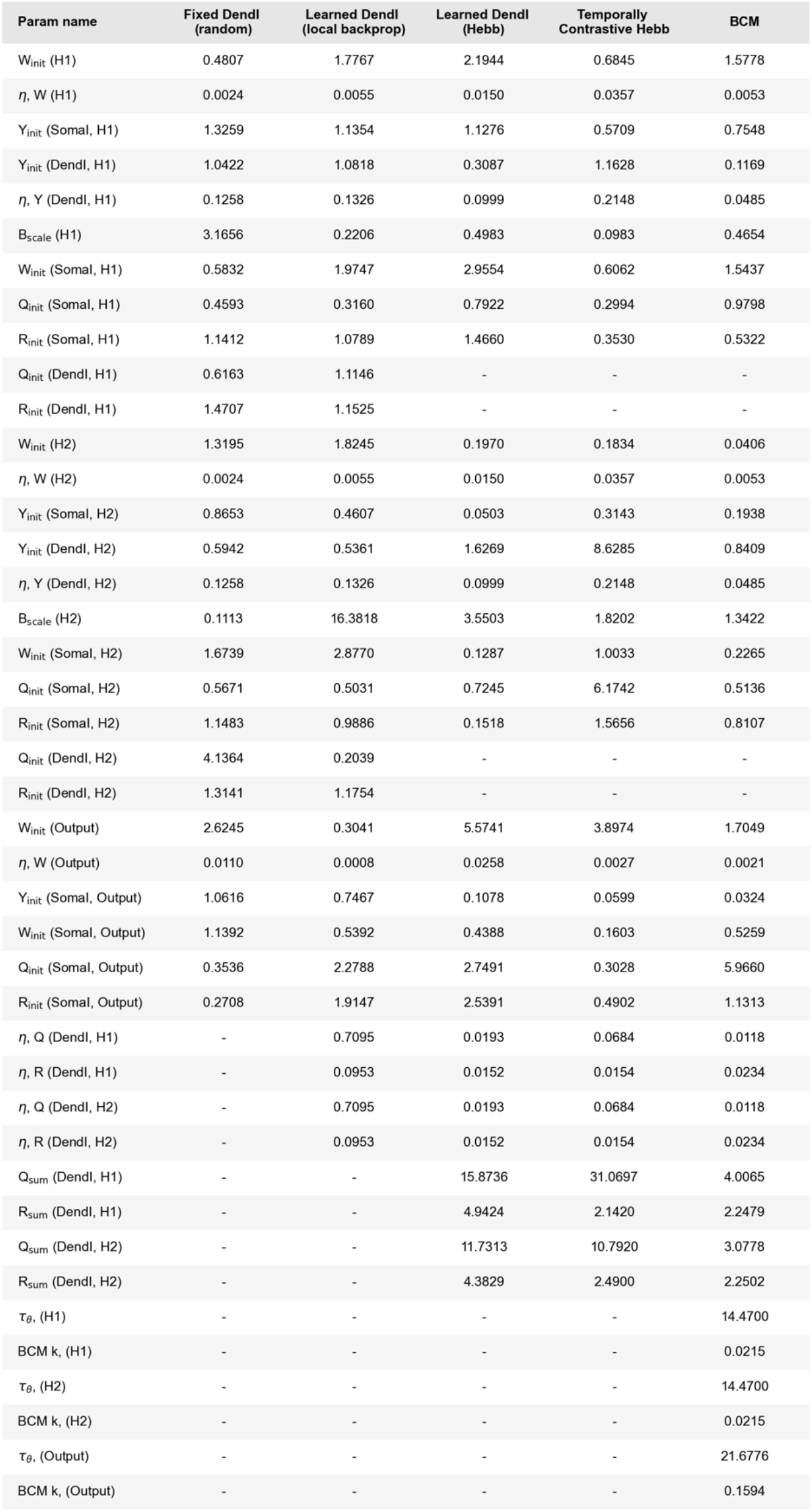
Hyperparameters for networks trained on handwritten digit classification (Part 2).

**Supplementary Table S5.** Hyperparameters for networks trained on handwritten digit classification (Part 3).

| Param name | BTSP | LDS,<br>fixed top-down | LDS,<br>learned top-down | BTSP,<br>fixed top-down | BTSP,<br>learned top-down |
| --- | --- | --- | --- | --- | --- |
| $W_{init}$ (H1) | 3.5237 | 1.5073 | 1.3345 | 3.9821 | 3.6173 |
| $\eta$, W (H1) | 0.0101 | 0.0171 | 0.0178 | 0.0025 | 0.0080 |
| BTSP $\lambda$, (H1) | 0.0162 | - | - | 0.0158 | 0.0368 |
| BTSP $W_{max}$, (H1) | 0.3520 | - | - | 0.3067 | 0.3848 |
| $Y_{init}$ (Somal, H1) | 2.1960 | 0.7136 | 1.1995 | 0.9458 | 0.9531 |
| $Y_{init}$ (Dendl, H1) | 1.4309 | 0.1059 | 0.9117 | 0.8045 | 0.6644 |
| $\eta$, Y (Dendl, H1) | 0.0540 | 0.0383 | 0.1806 | 0.0137 | 0.0320 |
| $B_{scale}$ (H1) | 0.1534 | - | - | - | - |
| $W_{init}$ (Somal, H1) | 2.2087 | 2.8738 | 0.9968 | 3.6431 | 2.9392 |
| $Q_{init}$ (Somal, H1) | 0.7036 | 0.8390 | 0.3449 | 0.3386 | 0.3796 |
| $R_{init}$ (Somal, H1) | 1.1848 | 1.2690 | 0.8168 | 0.8938 | 0.8945 |
| $\eta$, Q (Dendl, H1) | 0.1477 | 0.0859 | 0.0305 | 0.1545 | 0.0859 |
| $Q_{sum}$ (Dendl, H1) | 22.1510 | 9.1341 | 21.4930 | 32.7122 | 40.8871 |
| $\eta$, R (Dendl, H1) | 0.0517 | 0.0054 | 0.0584 | 0.0280 | 0.1953 |
| $R_{sum}$ (Dendl, H1) | 2.5624 | 0.0566 | 4.0984 | 1.6384 | 5.2394 |
| $W_{init}$ (H2) | 0.8324 | 1.3261 | 0.7726 | 1.3294 | 1.8581 |
| $\eta$, W (H2) | 0.0101 | 0.0171 | 0.0178 | 0.0025 | 0.0080 |
| BTSP $\lambda$, (H2) | 0.0162 | - | - | 0.0158 | 0.0368 |
| BTSP $W_{max}$, (H2) | 0.4408 | - | - | 0.3840 | 0.4818 |
| $Y_{init}$ (Somal, H2) | 0.2761 | 1.0827 | 0.5477 | 0.1948 | 0.2615 |
| $Y_{init}$ (Dendl, H2) | 6.2109 | 1.0116 | 1.1691 | 4.7234 | 3.7733 |
| $\eta$, Y (Dendl, H2) | 0.0540 | 0.0383 | 0.1806 | 0.0137 | 0.0320 |
| $B_{scale}$ (H2) | 5.9497 | - | - | - | - |
| $W_{init}$ (Somal, H2) | 2.4756 | 1.0048 | 0.9279 | 2.5163 | 1.3713 |
| $Q_{init}$ (Somal, H2) | 0.3042 | 0.0130 | 0.3694 | 0.4541 | 0.7330 |
| $R_{init}$ (Somal, H2) | 1.0033 | 0.7750 | 0.8262 | 1.1843 | 1.0274 |
| $\eta$, Q (Dendl, H2) | 0.1477 | 0.0859 | 0.0305 | 0.1545 | 0.0859 |
| $Q_{sum}$ (Dendl, H2) | 53.4596 | 17.2363 | 19.8207 | 49.7326 | 15.2392 |
| $\eta$, R (Dendl, H2) | 0.0517 | 0.0054 | 0.0584 | 0.0280 | 0.1953 |
| $R_{sum}$ (Dendl, H2) | 2.5428 | 3.5820 | 4.4111 | 2.9325 | 2.2955 |
| $W_{init}$ (Output) | 8.5974 | 0.4323 | 1.3278 | 12.6410 | 12.9520 |
| $\eta$, W (Output) | 0.0258 | 0.0222 | 0.0162 | 0.0291 | 0.0274 |
| BTSP $\lambda$, (Output) | 0.0162 | - | - | 0.0158 | 0.0368 |
| BTSP $W_{max}$, (Output) | 0.4353 | - | - | 0.5854 | 0.7402 |
| $Y_{init}$ (Somal, Output) | 0.1893 | 0.2388 | 0.5080 | 0.2754 | 0.3090 |
| $W_{init}$ (Somal, Output) | 1.1341 | 0.3644 | 0.2541 | 1.3436 | 0.8117 |
| $Q_{init}$ (Somal, Output) | 0.1276 | 0.5277 | 0.0414 | 0.2703 | 1.6620 |
| $R_{init}$ (Somal, Output) | 1.0732 | 0.5435 | 0.4426 | 2.9279 | 2.5709 |
| $B_{init}$ (H1) | - | 0.0959 | - | 0.0133 | - |
| $B_{init}$ (H2) | - | 1.1218 | - | 2.7066 | - |
| $\eta$, B (H1) | - | - | 0.0245 | - | 0.0216 |
| $B_{sum}$ (H1) | - | - | 0.3539 | - | 1.0467 |
| $\eta$, B (H2) | - | - | 0.0336 | - | 0.0088 |
| $B_{sum}$ (H2) | - | - | 0.4448 | - | 0.5107 |

**Supplementary Table S6.** Hyperparameters for networks trained on two-dimensional spiral pattern classification.

| Param name | Feedforward ANN<br>(no hidden) | Feedforward ANN<br>(learned bias) | Feedforward ANN<br>(no bias) | Backprop (EIANN) | Dendritic<br>Target Propagation |
| --- | --- | --- | --- | --- | --- |
| $W_{init}$ (Output) | 5.3865 | 0.5434 | 3.5188 | 5.9778 | 5.4269 |
| $\eta$, $W$ (Output) | 0.0816 | 0.0108 | 0.0013 | 0.6722 | 0.0977 |
| $W_{init}$ (H1) | - | 3.0922 | 0.7662 | 2.3272 | 3.7442 |
| $\eta$, $W$ (H1) | - | 0.2344 | 0.0087 | 0.1770 | 0.0856 |
| $W_{init}$ (H2) | - | 1.0685 | 0.1340 | 5.1834 | 3.3963 |
| $\eta$, $W$ (H2) | - | 0.2344 | 0.0087 | 0.1770 | 0.0856 |
| $Y_{init}$ (Somal, H1) | - | - | - | 4.7081 | 1.7171 |
| $W_{init}$ (Somal, H1) | - | - | - | 1.0646 | 8.5268 |
| $Q_{init}$ (Somal, H1) | - | - | - | 0.7309 | 2.5573 |
| $R_{init}$ (Somal, H1) | - | - | - | 1.9856 | 3.8213 |
| $Y_{init}$ (Somal, H2) | - | - | - | 3.2740 | 1.9616 |
| $W_{init}$ (Somal, H2) | - | - | - | 3.3866 | 3.3568 |
| $Q_{init}$ (Somal, H2) | - | - | - | 0.6920 | 2.6728 |
| $R_{init}$ (Somal, H2) | - | - | - | 1.6564 | 2.5278 |
| $Y_{init}$ (Somal, Output) | - | - | - | 1.7630 | 1.1147 |
| $W_{init}$ (Somal, Output) | - | - | - | 2.2899 | 1.9824 |
| $Q_{init}$ (Somal, Output) | - | - | - | 1.4600 | 0.9623 |
| $R_{init}$ (Somal, Output) | - | - | - | 1.5592 | 3.9245 |
| $Y_{init}$ (Dendl, H1) | - | - | - | - | 0.7285 |
| $\eta$, $Y$ (Dendl, H1) | - | - | - | - | 0.2102 |
| $B_{scale}$ (H1) | - | - | - | - | 0.1879 |
| $\eta$, $Q$ (Dendl, H1) | - | - | - | - | 0.0803 |
| $Q_{sum}$ (Dendl, H1) | - | - | - | - | 9.4193 |
| $\eta$, $R$ (Dendl, H1) | - | - | - | - | 0.0134 |
| $R_{sum}$ (Dendl, H1) | - | - | - | - | 1.1712 |
| $Y_{init}$ (Dendl, H2) | - | - | - | - | 2.7135 |
| $\eta$, $Y$ (Dendl, H2) | - | - | - | - | 0.2102 |
| $B_{scale}$ (H2) | - | - | - | - | 0.7150 |
| $\eta$, $Q$ (Dendl, H2) | - | - | - | - | 0.0803 |
| $Q_{sum}$ (Dendl, H2) | - | - | - | - | 11.6900 |
| $\eta$, $R$ (Dendl, H2) | - | - | - | - | 0.0134 |
| $R_{sum}$ (Dendl, H2) | - | - | - | - | 2.1474 |

